# Resting state brain network segregation is associated with walking speed and working memory in older adults

**DOI:** 10.1101/2024.05.07.592861

**Authors:** Sumire D. Sato, Valay A. Shah, Tyler Fettrow, Kristina G. Hall, Grant D. Tays, Erta Cenko, Arkaprava Roy, David J. Clark, Daniel P. Ferris, Chris J. Hass, Todd M. Manini, Rachael D. Seidler

**Affiliations:** Department of Applied Kinesiology and Physiology, University of Florida, Gainesville, FL, USA; NASA Langley Research Center, Hampton, VA, USA; Department of Epidemiology, College of Public Health and Health Professions, and College of Medicine, University of Florida, Gainesville, FL, USA; Department of Biostatistics, University of Florida, Gainesville, FL, USA; Department of Neurology, University of Florida, Gainesville, FL, USA; Brain Rehabilitation Research Center, Malcom Randall VA Medical Center, Gainesville, FL, USA; J. Crayton Pruitt Family Department of Biomedical Engineering, University of Florida, Gainesville, FL, USA; Department of Health Outcomes and Biomedical Informatics, University of Florida, Gainesville, FL, USA

**Keywords:** Functional connectivity, resting state, aging, fMRI, behavior, segregation

## Abstract

Older adults exhibit larger individual differences in walking ability and cognitive function than young adults. Characterizing intrinsic brain connectivity differences in older adults across a wide walking performance spectrum may provide insight into the mechanisms of functional decline in some older adults and resilience in others. Thus, the objectives of this study were to: (1) determine whether young adults and high– and low-functioning older adults show group differences in brain network segregation, and (2) determine whether network segregation is associated with working memory and walking function in these groups. The analysis included 21 young adults and 81 older adults. Older adults were further categorized according to their physical function using a standardized assessment; 54 older adults had low physical function while 27 were considered high functioning. Structural and functional resting state magnetic resonance images were collected using a Siemens Prisma 3T scanner. Working memory was assessed with the NIH Toolbox list sorting test. Walking speed was assessed with a 400 m-walk test at participants’ self-selected speed. We found that network segregation in mobility-related networks (sensorimotor, vestibular) was higher in older adults with higher physical function compared to older adults with lower physical function. There were no group differences in laterality effects on network segregation. We found multivariate associations between working memory and walking speed with network segregation scores. The interaction of left sensorimotor network segregation and age groups was associated with higher working memory function. Higher left sensorimotor, left vestibular, right anterior cingulate cortex, and interaction of left anterior cingulate cortex network segregation and age groups were associated with faster walking speed. These results are unique and significant because they demonstrate higher network segregation is largely related to higher physical function and not age alone.

**Highlights:**

- Older adults with high and low physical function have network segregation differences.
- Network segregation for high-functioning older adults was more similar to that of young adults.
- Laterality of functional network segregation is not different between age groups.
- Higher network segregation is associated with faster walking speed.

## 1. INTRODUCTION

In 2015, persons aged 65 years or older represented ∼ 9% of the world population; by 2050 this number is expected to increase to ∼17% (He et al., 2016). Many adults over the age of 65 have difficulty walking, which can decrease their quality of life and result in a variety of complications such as increased risk of cardiovascular disease and decreased bone health (Inouye et al., 2007; Shafrin et al., 2017). However, some older adults (i.e., “super agers” or “master athletes”) maintain their function and outperform their age-matched peers (Geard et al., 2017; Klinedinst et al., 2023). Characterizing differences in brain function across a wide performance spectrum may provide insight into the mechanisms of walking decline in some older adults and resilience in others.

Measures of how functional networks in the brain are connected at rest can inform us about intrinsic brain characteristics, including the capacity for communication within and across networks. Resting-state functional connectivity is quantified by fluctuations in the BOLD signal during rest (i.e., with participants lying in the magnetic resonance (MR) scanner without performing a task) (Biswal et al., 1995). By using graph theory approaches, we can model the brain as nodes (regions of interest; ROIs) and edges (connections identified by correlation of fluctuations in the BOLD signal across different brain regions). This approach has allowed the identification of networks of nodes that are functionally but not necessarily monosynaptically connected.

Neural de-differentiation (or more diffuse brain activity) has been observed in task-based functional neuroimaging studies of older adults (e.g., greater bilateral prefrontal activation during cognitive tasks) (Cabeza, 2002; Park et al., 2004). In healthy young adults, there is a balance between within– and between-network connectivity. In typical aging, functional brain networks are less connected within-network, but more connected between networks, leading to less segregated functional networks (Chan et al., 2014; Deery et al., 2023). This example of age-related neural dedifferentiation describes decreased functional specificity of localized brain regions and broader connectivity between-networks (Koen et al., 2020; Li and Lindenberger, 1999).

Importantly, lower sensorimotor network segregation is associated with lower sensorimotor function in older adults (Cassady et al., 2020; Cassady et al., 2019). Reduced network segregation reflects more overlapping or interacting processes in the aging brain, which is potentially less efficient. A better understanding of associations between physical and cognitive function with age differences in network segregation is important, and may help identify therapeutic target(s) for age-related neurological impairment. Less segregated networks in Alzheimer’s disease and mild cognitive impairment are associated with poorer cognitive function and greater tau burden (Ewers et al., 2021; Fu et al., 2022; Iordan et al., 2022; Steward et al., 2023). In addition, higher network segregation is predictive of greater intervention-related cognitive gains (Cohen and D’Esposito, 2016; Gallen et al., 2016; Gallen and D’Esposito, 2019).

In this study we specifically assessed working memory performance, since working memory is a key function driving various cognitive behaviors (Dang et al., 2014; Engle et al., 1999; Kane et al., 2004). We also assessed walking speed; walking speed is associated with overall health, independence level, fall risk, cognition, and quality of life in community-dwelling adults (Abellan van Kan et al., 2009; Kim et al., 2016; Middleton et al., 2015). Both working memory and walking speed decline in aging (albeit at different speeds and magnitudes). Dorsolateral prefrontal cortex is a key area involved in working memory (Barbey et al., 2013). In addition, anterior cingulate cortex helps allocate resources to relevant tasks and filtering out attention for efficient and focused working memory (Lenartowicz and McIntosh, 2005; Seamans et al., 2024).

Walking performance relies on several areas of the brain such as sensorimotor cortex, vestibular areas, and visual and anterior cingulate cortex for navigation and error detection (Takakusaki, 2017; Wenderoth et al., 2005). In aging, walking requires more prefrontal resources (Chatterjee et al., 2019; Hawkins et al., 2018; Hwang et al., 2024), suggesting compensatory roles. Importantly, these behavioral functions are not isolated; for example, working memory can impact walking behavior by affecting the ability to plan and execute movement. Examining the role of the resting state brain network segregation in integrated behavioral functions may provide valuable insights into the mechanisms underlying cognitive and motor decline and potentially inform interventions to mitigate these effects.

In addition to reduced network segregation with age, the human brain also undergoes lateralization changes with aging (Sugiura, 2016). Reduced lateralization in functional brain activation has been shown during cognitive task performance for older adults. This finding has led to the development of the hemispheric asymmetry reduction in older adults (HAROLD) model, which proposes that aging is associated with a more bilateral distribution of brain activity (Cabeza, 2002). This model has also been supported in studies of motor control. Young adults generally show higher activation in the sensorimotor cortex contralateral to the moving limb. In contrast, there is increased brain activation in the hemisphere ipsilateral to the moving limb in older adults, resulting in brain activation that is more bilaterally symmetrical and diffuse (Bernard and Seidler, 2012; Seidler et al., 2010). Independent component analysis has demonstrated that resting-state functional connectivity in the sensorimotor networks has both left and right dominant components, but lateralization decreases with increasing age (Agcaoglu et al., 2015). This finding suggests that the age-related changes in lateralization are not limited to cognitive processing, but rather reflect a more general shift towards a more bilateral distribution of brain activity. The factors that contribute to age-related changes in lateralization are still not fully understood, but it is thought that both genetic and environmental factors play a role (Liu et al., 2009; Yoon et al., 2010). Age-related changes in resting state brain network segregation have been speculated to play a role in cognitive decline in older adults (Deery et al., 2023), but the role of hemisphere-specific network segregation on sensorimotor function has not been investigated. Investigating the hemispheric-specific role in cognitive and walking function has the potential to identify new neural mechanisms underlying mobility variability in older adults.

The objectives of this study were to: (1) determine whether young adults and older adults with high and low physical function show group differences in resting state brain network segregation, and (2) determine whether resting state brain network segregation is associated with working memory and walking speed in these groups. Neural de-differentiation is associated with age-related decline in cognition (Koen et al., 2020; Li and Lindenberger, 1999; Li et al., 2001). We recruited young adults and older adults who were categorized into high and low function according to their performance on a standardized battery of physical tasks. We hypothesized that: (1a) older adults would demonstrate lower functional sensorimotor network segregation compared to younger adults, and that older adults with lower mobility function would demonstrate greater differences than older adults with higher mobility function when compared to healthy younger adults, (1b) older adults with lower mobility function would demonstrate less lateralized functional connectivity compared to older adults with higher mobility function and younger adults, and (2) higher brain network segregation would be associated with faster walking speed and better working memory performance in older adults. Specifically, we hypothesized that higher brain network segregation in the motor-related networks (sensorimotor, vestibular, and visual) will be associated with faster walking speed, and that the higher dorsolateral prefrontal cortex and anterior cingulate cortex network segregation will be associated to working memory performance.

## 2. MATERIAL AND METHODS

### 2.1. Participants

This study enrolled community-dwelling volunteers with 23 young adults, and 87 older adults. Participants recruited for this study were part of a larger NIH U01 study at the University of Florida (U01AG061389). Briefly, all participants were required: to be able to walk 400 m within 15 minutes without sitting or assistance; be English speaking; to have no significant medical event within the past six months; to have no major neurological injuries (e.g., stroke, vestibular dysfunction, traumatic brain injury, dementia); and to meet MR imaging eligibility (Full exclusion and inclusion criteria outlined: Clark et al., 2019). Older adults were further categorized into their physical function level, based on their Short Physical Performance Battery score. The battery included a short distance usual walk speed test, balance test and chair stand speed. Each test was graded according to normative values and tallied for a total score that ranged from 0-12 with 12 serving as the highest physical function (Guralnik et al., 1994). The study screened in those who scored > 10 for the high functioning (n = 27) group and those who scored ≤ 10 for the low functioning (n = 54) group. The sample size for the older adults with lower physical function was the largest because it was the target sample of interest; in most studies of human aging, the older adult cohort is relatively fit due to inclusion/exclusion criteria (Brach et al., 2023). Therefore, we especially wanted to study the variability in resting state brain mechanisms in older adults with lower physical function. As mentioned earlier, this is part of a larger study (Clark et al., 2019) which aims to investigate longitudinal changes in brain -behavior relationships, especially in the lower functioning older adults. The study design of the larger study led to an uneven group size. In addition, the sample size between older adults with lower function and higher function came out unequal because our data collection was disrupted due to the COVID-19 pandemic. Group characteristics for participants who were included in the final analysis of this study are presented in Table 1. Participants who identified as left-handed were excluded from the analyses to investigate laterality (2 younger adults, 2 older adults with higher physical function, and 4 older adults with lower physical function were excluded; participants included in final analysis included 21 younger adults, 27 older adults with higher physical function, and 54 older adults with lower physical function). Participants’ written informed consent was obtained prior to the study, which was approved by the Institutional Review Board of the University of Florida, Gainesville, FL, USA (Protocol# 201802227).

**Table 1.**
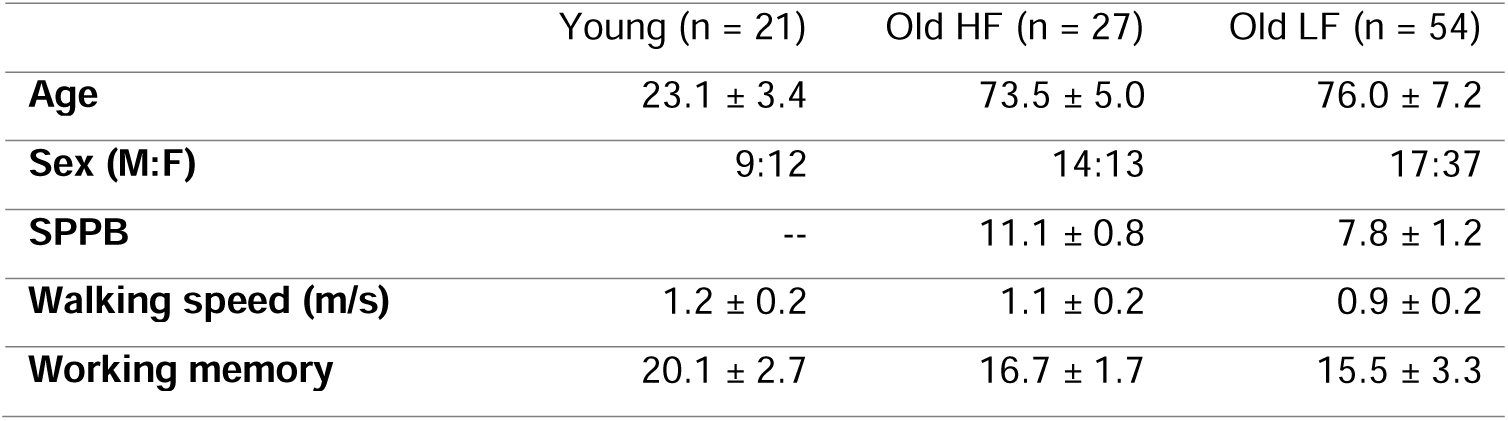
Participant characteristics. HF = Older adults with high physical function. LF = Older adults with low physical function. There was one young adult and one older adult in the higher function group with missing gait speed. For working memory, 3 young adults, 4 older adults with high physical function, and 19 older adults with lower physical function were excluded due to missing data. SPPB = Short Physical Performance Battery (Max score = 12; Higher score = higher physical function). Working memory was assessed with the list sorting test on the NIH Toolbox (Max score = 26; Higher score = higher working memory).

### 2.2. Brain Imaging

Brain images were collected using a Siemens Prisma 3T scanner (Siemens Healthcare, Erlangen, Germany) with a 64-channel head coil. Structural T1-weighted images were collected using a magnetization-prepared rapid gradient echo (MPRAGE) sequence with the following parameters: repetition time (TR) = 2000 ms, echo time (TE) = 2.99 ms, flip angle = 8°, FOV = 167 mm × 256 mm × 256 mm, voxel size = 0.80 mm^3^. Whole-brain resting-state functional MR images were collected using a multiband, interleaved echo planar imaging (EPI-BOLD) sequence with the following parameters: TR = 1500 ms, TE= 30 ms, flip angle= 70°, FOV= 240 x 240 x 165 mm, matrix=64×64, voxel size= 2.5 mm^3^, 66 axial slices, slice thickness = 2.5 mm, acceleration factor = 3, duration = 8 minutes.

### 2.3. Neuroimaging preprocessing

Neuroimaging data analyses were performed using the Statistical Parametric Mapping 12 software (SPM12; www.fil.ion.ucl.ac.uk/spm) and the CONN toolbox version 19 (https://www.nitrc.org/projects/conn) (Whitfield-Gabrieli and Nieto-Castanon, 2012) in MATLAB 2020b (Mathworks, Natick, MA, USA). Preprocessing steps included: slice-time correction, realignment and unwarping, movement correction, bias correction, co-registration of functional and anatomical images and normalization to a Montreal Neurological Institute 152 (MNI152) template using Advanced Normalization Tools (ANTs, https://github.com/ANTsX/ANTs) (Avants et al., 2011), then spatial smoothing (Gaussian smoothing kernel 4mm full width at half maximum). We used the Artifact Detection Tool (ART, www.nitrc.org/projects/artifact_detect/) to detect and correct for movement artifacts. Within a run, a volume was considered an outlier and covaried out if the participant’s composite movement was equal to or greater than 0.5 mm and a global mean intensity threshold at 5 standard deviations of the mean image intensity (Mean number of outliers per group: Young adults = 0.38 ± 1.36; Older adults with high physical function= 0.19 ± 0.96; Older adults with lower physical function = 0.52 ± 1.71; ANOVA showed no significant differences: F(2,99) = 0.46, p = 0.633).

Additional denoising was done with CONN; we estimated confounding noise arising from non-neuronal sources using the anatomical component-based noise correction (aCompCor) tool (Behzadi et al., 2007). This method extracts signals from eroded white matter and CSF masks applied to unsmoothed resting state fMRI data. These signals underwent principal components analysis, with the top five components for white matter and CSF signals reflecting noise and physiological artifact (e.g., pulsation, respiration). Covariates modeling physiological noise sources were included as nuisance regressors during the denoising steps. Denoising was performed by regressing out the confounding effects of the following nuisance regressors: six head motion parameters (translations and rotations used during realignment) and their six first-order temporal derivatives, five principal components modeling physiological noise from within the white matter, five principal components modeling physiological noise from within the CSF, and single volume “scrubbing” regressors modeling outlier volumes that exceeded based on movement and global brain signal thresholds. Following denoising, we filtered the data using a temporal band-pass filter of 0.008-0.090 Hz to examine the slower frequency band of interest and to exclude higher frequency sources of noise (Biswal et al., 1995; Damoiseaux, 2017).

To improve cerebellar normalization to MNI space, the cerebellum from each anatomical image was segmented from the brain using the CERES (CEREbellum Segmentation) (Romero et al., 2017) pipeline. They were then normalized to the spatially unbiased infra-tentorial template (SUIT) cerebellar template (Diedrichsen, 2006).The SUIT cerebellar template shows greater anatomical detail than the whole brain MNI152 template, allowing for more accurate cerebellar normalization (Diedrichsen, 2006). We applied the CERES-generated cerebellar masks to the co-registered functional images, allowing extraction of the cerebellum. The extracted fMRI cerebellums were then normalized to the SUIT cerebellar template using the warp fields generated for the T1 scans.

### 2.4. Connectivity analysis

We used the CONN (v19b) toolbox (Whitfield-Gabrieli and Nieto-Castanon, 2012) for all connectivity analyses. For identification of our networks we used a seed-to-voxel analysis with seed regions of interest (ROIs). We derived the seed ROI coordinates from previous findings that identified brain regions to be associated with mobility (Left and right sensorimotor (Zhao et al., 2019), left and right vestibular (zu Eulenburg et al., 2012), default mode (Schmidt et al., 2017), and visual (Shen et al., 2019)). The left and right dorsolateral and left and right anterior cingulate cortex seed ROIs were identified using custom ROIs for this study. The seed regions were identified using the Neurosynth meta-analysis with “working memory” search terms for the dorsolateral prefrontal cortex ROI and “anterior cingulate” for the anterior cingulate cortex ROI (Yarkoni et al., 2011); both seed regions were anatomically constrained to the gray matter of dorsolateral prefrontal cortex and anterior cingulate cortex, respectively. We had a priori hypotheses regarding the dorsolateral prefrontal cortex and the anterior cingulate cortex in our larger study that dorosolateral prefrontal and anterior cingulate regions will be associated with age-related functional decline (Clark et al., 2019).

Therefore, we wanted to identify brain networks associated with each of these regions of interest and also preserve the typical brain networks used in other whole brain network analyses (e.g., Gordon et al., 2016; Power et al., 2011). To define our network nodes we performed the seed-to-voxel analyses on 110 subjects from the Washington University 120 dataset (data were obtained from the OpenfMRI database, accession number: ds00243). We used an independent, open-source dataset to define our functional network nodes to increase generalizability for future studies, and to avoid circular reasoning (Kriegeskorte et al., 2009) and bias by the specific characteristics of the dataset used for the results of the manuscript.

First seed-to-voxel analysis was performed with the MNI normalized whole brain images as the target (with the cerebellum masked), then a second seed-to-voxel analysis was performed with the SUIT normalized cerebellum images as the target. For each seed-to-voxel analysis performed, we implemented a second level analysis using the default statistical thresholds for cluster-based inferences (Worsley et al., 1996). The resulting T map images with a-values over 12, and local maxima separated by 8 mm were identified. A 4 mm radius sphere was created around each cluster. We combined the cerebellar coordinates with the whole brain coordinates for each respective seed. In total, we identified 279 nodes in 10 networks for analysis in this study: (1) Left sensorimotor network (26 nodes), (2) right sensorimotor network (22 nodes), (3) left vestibular network (26 nodes), (4) right vestibular network (26 nodes), (5) left dorsolateral prefrontal cortex network (21 nodes), (6) right dorsolateral prefrontal cortex network (22 nodes), (7) left anterior cingulate cortex network (27 nodes), (8) right anterior cingulate cortex network (30 nodes), (9) default mode network (48 nodes), and (10) visual network (31 nodes), (Figure 1A; MNI coordinates for the ROIs are in Supplementary table 1).

**Figure 1.**
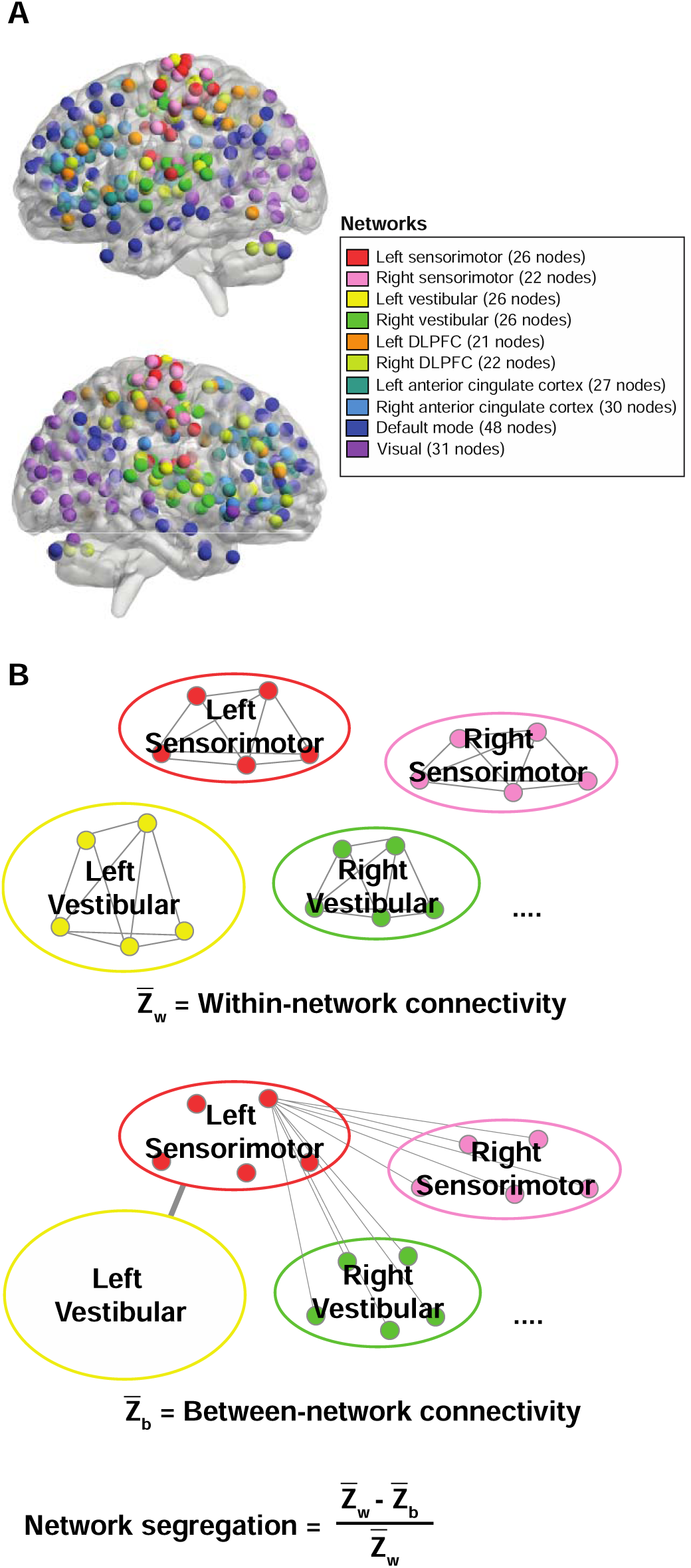
Network segregation analysis. A. Network connectivity nodes included in the study. In total, 279 nodes in 10 networks were identified for analysis. BrainNet Viewer (https://www.nitrc.org/projects/bnv/; Xie et al., 2013) was used to illustrate the nodes. B. For each network, mean within-network connectivity was calculated as the mean z-values between the nodes within the network (i.e., Left sensorimotor within-network connectivity = mean z-value of all nodes in the network to all other nodes in the left sensorimotor network). Between-network connectivity was calculated as the mean of z-values between each node in the network and all the nodes in other networks (i.e., Left sensorimotor between-network connectivity = mean z-value between all nodes in left sensorimotor network and all other nodes in the brain). Network segregation score for each network was quantified as the difference of the mean within-network connectivity and the mean between-network connectivity divided by the mean within-network connectivity.

We applied a node-to-node first-level functional connectivity analysis to the current dataset using the 10 networks described above. Resting-state time series within each node were extracted from the unsmoothed functional images and the mean time series was computed for each node (averaged across voxels). Then, we computed a correlation analysis for all nodes, producing a 279 x 279 cross-correlation matrix for each participant. The correlation coefficients (i.e., graph edges) were converted into z-values using Fisher’s r-to-z transformation. Negative-weighted edges were set to zero in each participant’s correlation matrix to avoid potential misinterpretation of negative edge weights (Chan et al., 2014).

We quantified network segregation for each network as the difference of the mean within-network connectivity and the mean between-network connectivity divided by the mean within-network connectivity (Eq. 1, Figure 1B).

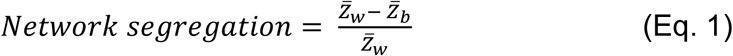

Within-network connectivity (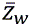) is the mean of Fisher z-transformed correlations between nodes within each network and between-network connectivity (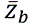) is the mean of Fisher z-transformed correlations between nodes of one network to all nodes in all other networks (Chan et al., 2014).

### 2.5. Walking function

To measure walking speed (m/s), we conducted a 400 m-walk test where participants were asked to walk around a pair of traffic cones set 20 meters apart. We instructed participants to complete 10 laps (each lap being 40 meters long) by walking at their preferred pace. Participants could stop for up to 1 minute if they were feeling fatigued.

### 2.6. Cognitive Function

We used a subset of tests from the NIH toolbox cognitive battery to assess cognitive function as part of the larger study. The list sorting test assessed working memory capacity and recall, presenting the participants with a series of visual and auditory stimuli using an iPad (Apple Inc., Cupertino, CA) (Tulsky et al., 2014). Participants were asked to recall and sequence these stimuli in a specific order. Each visual stimulus was displayed on the iPad screen for two seconds followed by an audible stimulus of the word (e.g., an image of a horse was shown on the screen while a computerized voice read the word “horse”). We instructed participants to remember each stimulus, arrange them mentally by size, and then verbally list them in that order. We used the raw score for the working memory test, which ranges from 0 to 26. These scores were chosen over age-corrected standard scores to assess the participant’s absolute performance in each test, as opposed to their performance relative to the NIH Toolbox normative sample. This approach to interpreting computed and raw scores aligns with the NIH Toolbox Scoring and Interpretation guide’s recommendations.

### 2.7. Statistical analysis

Group differences in segregation scores, between-network, and within-network connectivity were assessed through one-way ANCOVA, with biological sex as a covariate. Post-hoc pairwise comparisons were conducted with Bonferroni-Holm corrections.

The relationship between functional measures and network segregation scores was assessed with a canonical correlation analysis. Canonical correlation analysis is a multivariate analytical approach that measures the relationship between two datasets. We identified a “Function variable dataset” with walking speed, working memory, age/physical function group, and biological sex, and a “Network segregation dataset” with network segregation scores from the 10 identified networks. Canonical correlation analysis was conducted twice: first with the age groups (Older adult = -1 | Younger adult = 1), second with the physical function groups, with the younger adults omitted from the analysis (Lower function = –1 | Higher function = 1). We chose to do canonical correlation analysis twice with a maximum of two groups in our categorical variable to maintain simplicity and transparency in the inference. All variables were standardized before canonical correlation analysis to make the canonical coefficients comparable across the variables. Canonical correlations range from 0 to 1, 0 being no relationship and 1 being a perfect relationship between the two datasets.

As a post-hoc analysis, we tested two multiple linear regression models with age groups: 1. Walking speed and 2. Working memory, both with biological sex covaried out. Similarly, for functional groups we tested multiple linear regression models with older adults only. Due to missing data, one young adult and one older adult with higher physical function were excluded from the linear regression with walking speed, and 3 young adults, 4 older adults with high physical function, and 19 older adults with lower physical function were excluded from the regressions with working memory scores.

Univariate linear regressions between the dependent variable and segregation scores and age groups for each network determined the variables and network-age group interactions that were considered in the step-wise multiple regression model. The univariate regression-based predictor selection mimics methods like sure independent screening (SIS) (Fan and Lv, 2008). Instead of using the magnitude of estimated coefficients, we consider p-values, which account for the standard error. Stepwise forward selection introduces bias due to the need for a pre-specified order of inclusion, while backward selection is problematic because of the limited sample size of our study. Therefore, we use backward selection only after screening a subset of variables with component-wise regressions.

All statistical significance was established with an alpha level = 0.05. We used RStudio (R version 4.3.0) for analysis. For canonical correlation analysis results, we applied the “CCA” package (González et al., 2008) which includes a regularization approach to handle missing data.

## 3. RESULTS

### 3.1. Segregation scores

There were significant group effects for the sensorimotor network (F(2, 98) = 6.54, p = 0.002), vestibular network (F(2, 98)= 19.78, p <0.001), dorsolateral prefrontal cortex (F(2,98)=3.45, p = 0.036), and anterior cingulate cortex networks (F(2,98)=3.87, p =0.024) (Figure 2, Table 2).

**Figure 2.**
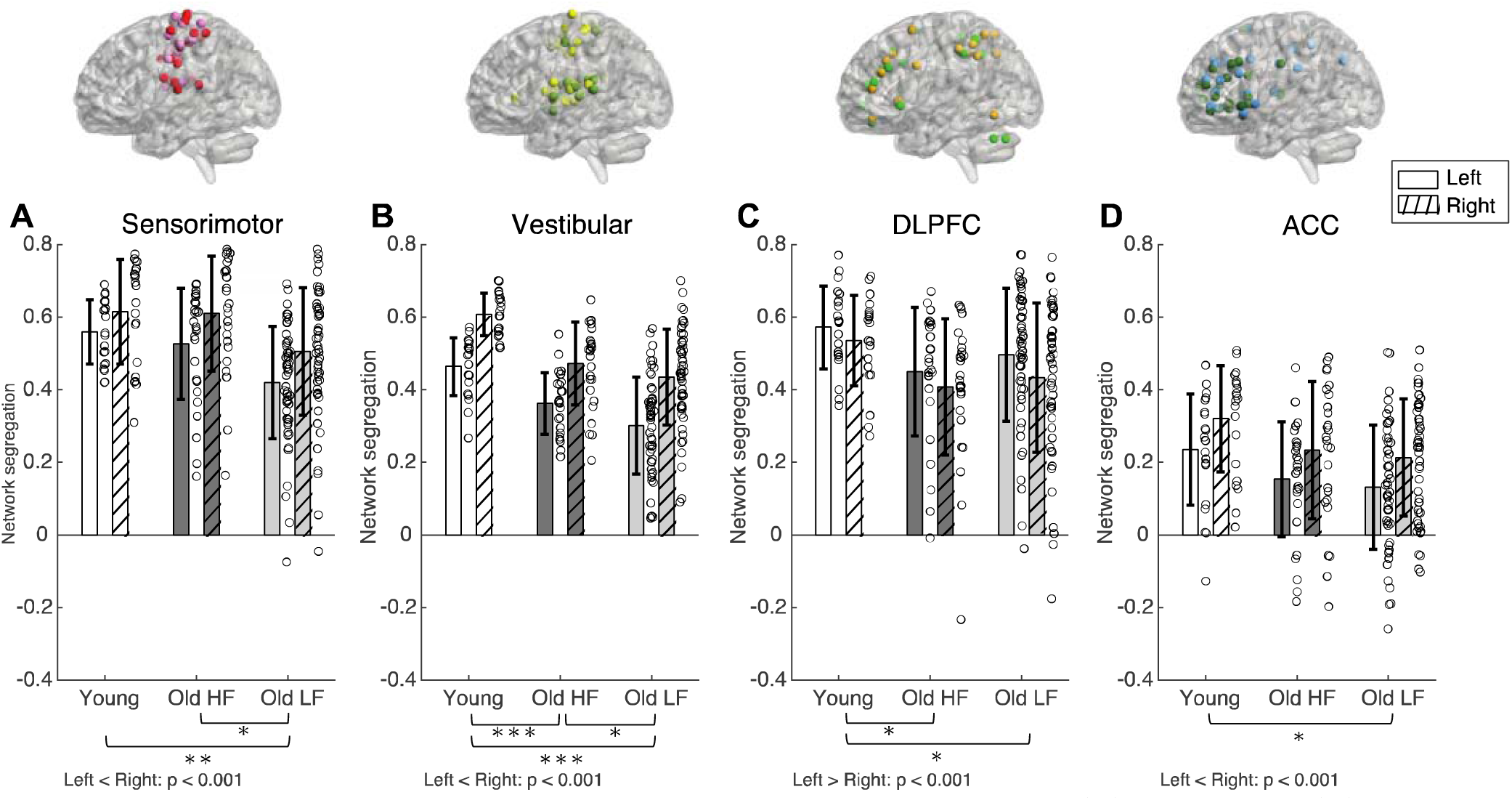
Network segregation score for left and right sensorimotor. (A), vestibular (B), dorsolateral prefrontal cortex, (C) and anterior cingulate cortex (D) networks. Brackets indicate statistically significant comparisons (p< 0.05) between groups from post-hoc tests. Color scheme for network nodes follow Figure 1. DLPFC = Dorsolateral Prefrontal Cortex. ACC = Anterior Cingulate Cortex. HF = High physical function. LF = Low physical function. * = p-value 0.010 ≤ p < 0.050. ** = p-value 0.001 ≤ p < 0.009. ***= p < 0.001

**Table 2.** Statistical output for post-hoc pairwise group comparisons in network segregation scores. P-values are adjusted for multiple comparisons using the Holm-Bonferroni method. Y = Young adults. OA HF = Older adults with high physical function. OA LF = Older adults with low physical function. DLPFC = Dorsolateral Prefrontal Cortex. ACC = Anterior Cingulate Cortex. DMN = Default Mode Network.

| Network | Contrast | Estimate | SE | t-ratio | p-value |
| --- | --- | --- | --- | --- | --- |
| **Sensorimotor** | Y - OA HF | 0.025 | 0.042 | 0.579 | 0.564 |
|  | Y - OA LF | 0.118 | 0.038 | 3.144 | 0.007 |
|  | OA LF - OA- HF | 0.094 | 0.035 | 2.686 | 0.017 |
| **Vestibular** | Y - OA HF | 0.118 | 0.030 | 3.912 | < 0.001 |
|  | Y - OA LF | 0.168 | 0.027 | 6.289 | < 0.001 |
|  | OA LF - OA- HF | 0.050 | 0.025 | 2.020 | 0.046 |
| **DLPFC** | Y - OA HF | 0.118 | 0.048 | 2.439 | 0.049 |
|  | Y - OA LF | 0.098 | 0.043 | 2.285 | 0.049 |
|  | OA LF - OA- HF | -0.020 | 0.040 | -0.506 | 0.614 |
| **ACC** | Y - OA HF | 0.081 | 0.044 | 1.845 | 0.136 |
|  | Y - OA LF | 0.109 | 0.039 | 2.779 | 0.020 |
|  | OA LF - OA- HF | 0.027 | 0.036 | 0.751 | 0.454 |
| **DMN** | Y - OA HF | 0.027 | 0.051 | 0.526 | 0.811 |
|  | Y - OA LF | 0.062 | 0.045 | 1.369 | 0.523 |
|  | OA LF - OA- HF | 0.035 | 0.042 | 0.836 | 0.811 |
| **Visual** | Y - OA HF | 0.103 | 0.042 | 2.449 | 0.048 |
|  | Y - OA LF | 0.083 | 0.037 | 2.227 | 0.057 |
|  | OA LF - OA- HF | -0.020 | 0.035 | -0.580 | 0.563 |

Sensorimotor network segregation was lower in older adults with lower physical function compared to younger adults and older adults with higher physical function, but sensorimotor network segregation was not significantly different between young adults and older adults with higher physical function. The left sensorimotor network was less segregated compared to the right (F(1, 98) = 55.67, p < 0.001); there were no Group x Laterality interaction effects (F(2,98) = 0.84, p = 0.437). Vestibular network segregation was higher in young adults compared to both older adult groups with higher and lower physical function. Older adults with higher physical function had higher vestibular network segregation compared to older adults with lower physical function. Left vestibular network was less segregated compared to the right (F(1,98) = 141.86, p < 0.001), but there were no Group x Laterality interaction effects (F(2,98) = 0.60, p =0.553).

Dorsolateral prefrontal cortex network segregation was higher in young adults compared to both older adult groups with higher and lower physical function, but not different between older adult sub-groups. In contrast to the sensorimotor and vestibular networks, dorsolateral prefrontal cortical network was more segregated in the left compared to the right (F(1,98) = 15.79, p < 0.001). Anterior cingulate cortex network segregation was higher in young adults compared to older adults with lower physical function, but not significantly different compared to older adults with higher physical function and between older adult sub-groups. Anterior cingulate cortex network was less segregated in the left compared to the right (F(1,98) = 31.28, p < 0.001). Group x Laterality interaction effects were not statistically significant for these two networks (Dorsolateral prefrontal cortex: F(2,98) = 0.69, p = 0.504; Anterior cingulate cortex: F(2,98) = 0.02, p =0.983)

We included the visual network in the analyses because of its role during walking navigation, and we included the default mode network as a ‘control’ network (we did not expect that default mode network segregation would be associated with working memory performance or walking speed). Since these two networks have medial seed regions, they do not contain laterality outcomes; thus, we present them separately in Figure 3. The visual network was more segregated in younger adults compared to older adults with high physical function, but not significantly different compared to older adults with low physical function and between older adult sub-groups (Group effect: F(2,98)=3.39, p = 0.038). Default mode network connectivity was not significantly different between groups (F(2,98)=1.03, p = 0.361).

**Figure 3.**
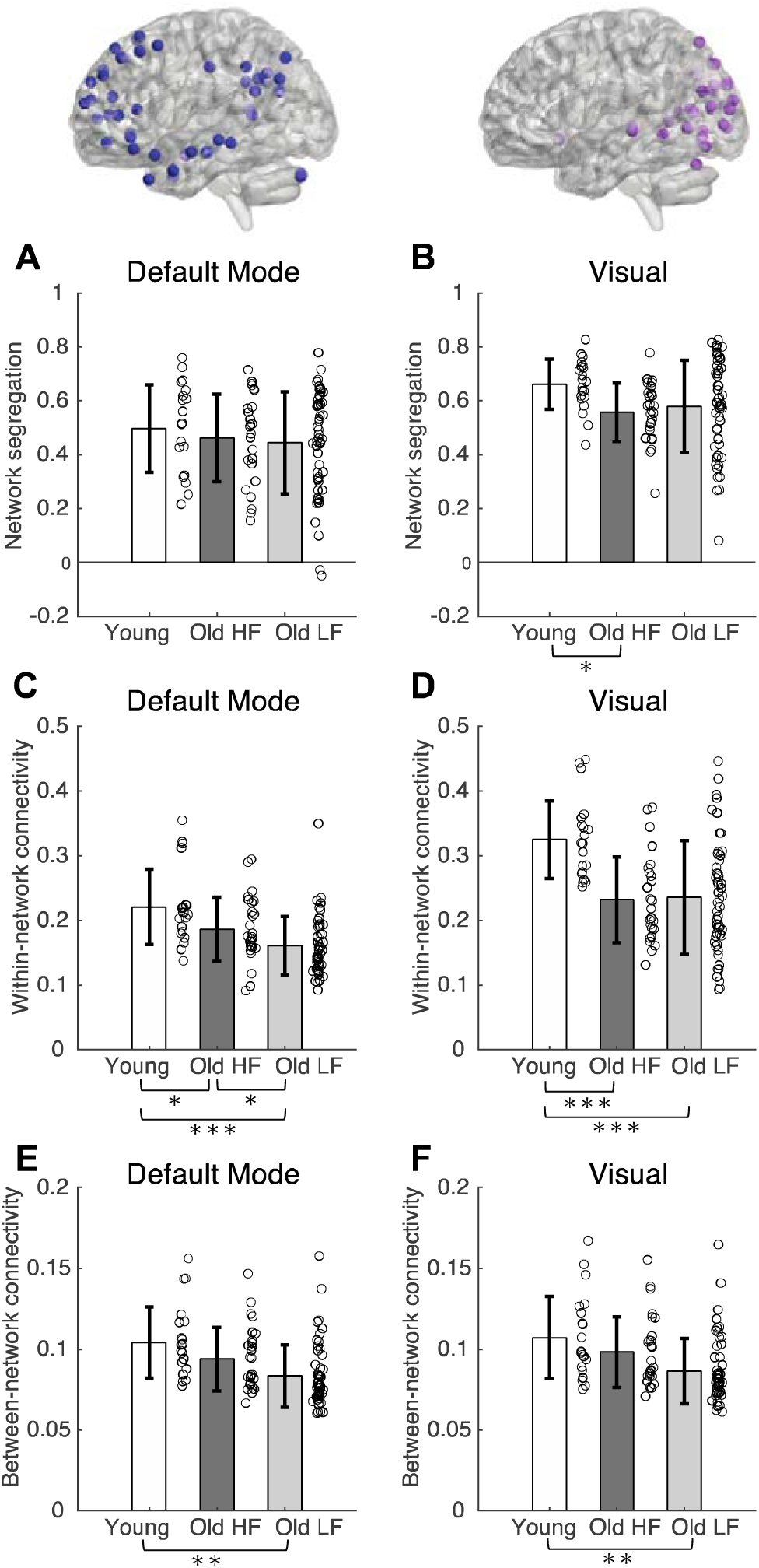
Network segregation (A-B), within-network connectivity (C-D) and between-network connectivity (E-F) for default mode (A, C, E), visual (B, D, F) networks. Brackets indicate statistically significant comparisons (p< 0.05) between groups from post-hoc tests. Color scheme for network nodes follow Figure 1. DLPFC = Dorsolateral Prefrontal Cortex. ACC = Anterior Cingulate Cortex. HF = High physical function. LF = Low physical function. * = p-value 0.010 ≤ p < 0.050. ** = p-value 0.001 ≤ p < 0.009. ***= p < 0.001

### 3.2. Within– and between-network connectivity

To parse out the root of segregation score differences, we also examined group and laterality differences in within-network connectivity and between-network connectivity, both of which make up the network segregation scores (Figure 3 C-F and Figure 4, Tables 3 and 4). In general, within-network connectivity was higher in younger adults compared to older adults. Sensorimotor within-network connectivity was lower in older adults with lower physical function compared to younger adults and older adults with higher physical function, but was not significantly different between younger adults and older adults with higher physical function (Group effect: F(2,98) = 10.24, p<0.001). Left and right sensorimotor within-network connectivity was not significantly different (F(1,98) = 2.50, p = 0.117). Vestibular and anterior cingulate cortex within-network connectivity was significantly different between all groups (Vestibular: F(2,98) = 22.01, p<0.001; Anterior cingulate cortex: F(2, 98) = 19.72, p<0.001); within-network connectivity was higher in younger adults compared to both older adult groups, and within-network connectivity was higher in older adults with higher physical function compared to older adults with lower physical function. For both the vestibular and anterior cingulate cortex networks, within-network connectivity was higher on the right side compared to the left side (Laterality effect, Vestibular: F(1,98)=57.70, p < 0.001; Laterality effect, Anterior cingulate cortex: F(1,98) = 37.99, p < 0.001). Dorsolateral prefrontal cortex within-network connectivity was higher in young adults compared to both older adult groups, but dorsolateral prefrontal cortex network within-network connectivity was not significantly different between older adults with higher and lower physical function (Group effect: F(2, 98) = 17.26, p < 0.001). Dorsolateral prefrontal cortex within-network connectivity was higher on the left side compared to the right side (F(1,98) = 28.47, p < 0.001).

**Figure 4.**
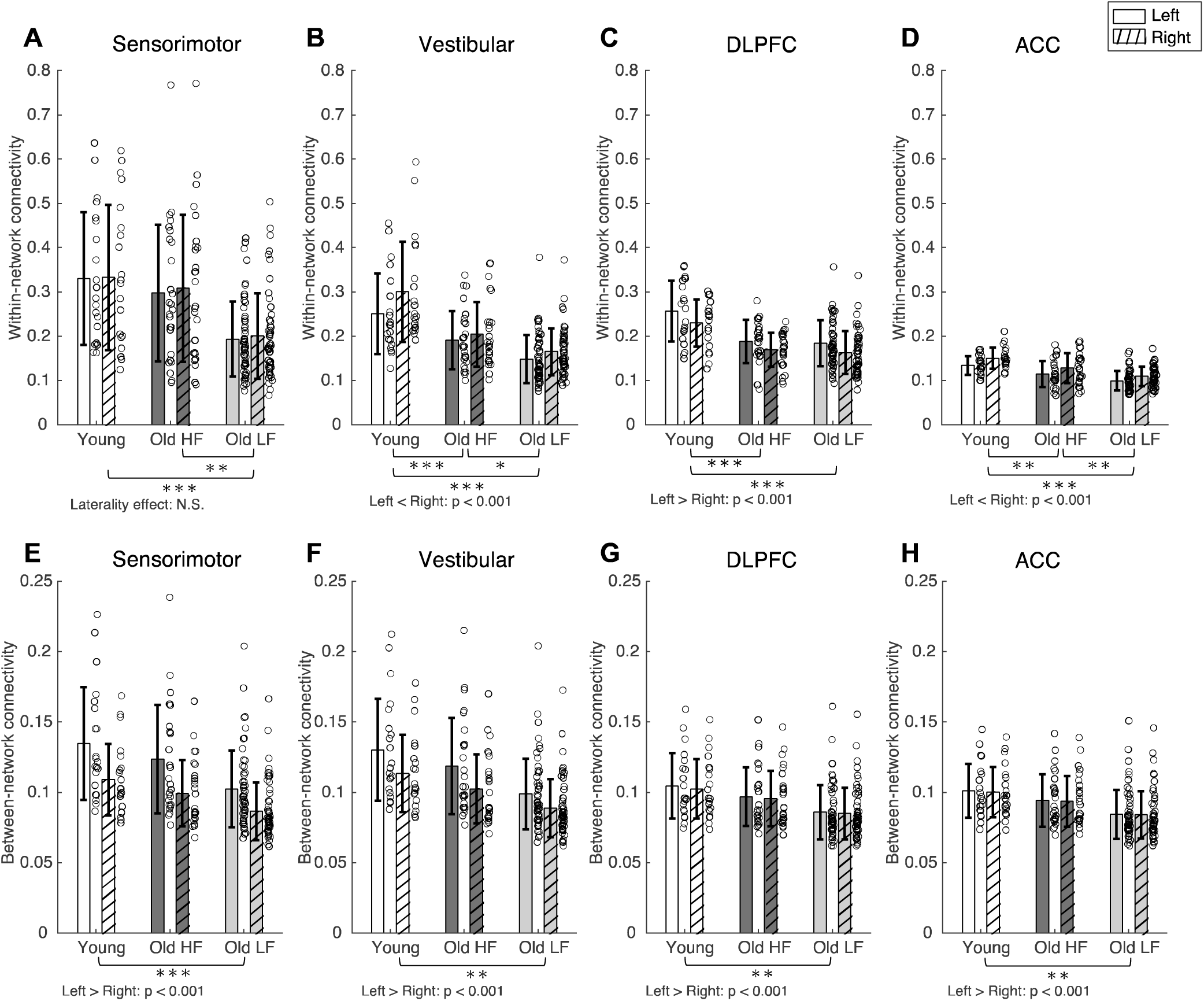
Within-network connectivity (A-D) and between-network connectivity (E-H) for sensorimotor (A, E), vestibular (B, F), dorsolateral prefrontal cortex (C, G), and anterior cingulate cortex (D, H) networks. Brackets indicate statistically significant comparisons (p< 0.05) between groups from post-hoc tests. DLPFC = Dorsolateral Prefrontal Cortex. ACC = Anterior Cingulate Cortex. HF = High physical function. LF = Low physical function. * = p-value 0.010 ≤ p < 0.050. ** = p-value 0.001 ≤ p < 0.009.

**Table 3.**
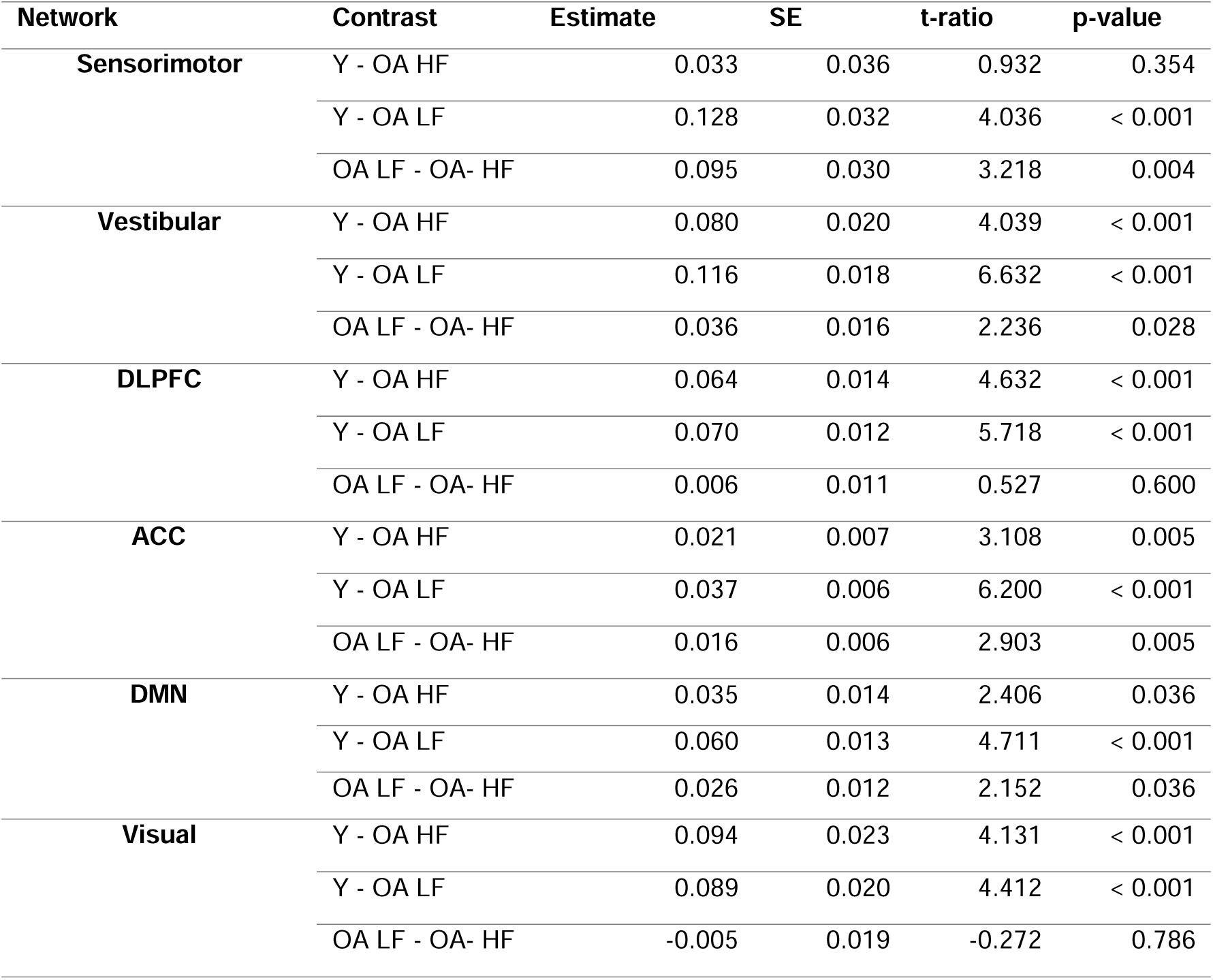
Statistical output for post-hoc pairwise group comparisons in within-network connectivity scores. P-values are adjusted for multiple comparisons using the Holm-Bonferroni method. Y = Young adults. OA HF = Older adults with high physical function. OA-LF = Older adults with low physical function. DLPFC = Dorsolateral Prefrontal Cortex. ACC = Anterior Cingulate Cortex. DMN = Default Mode Network.

| Network | Contrast | Estimate | SE | t-ratio | p-value |
| --- | --- | --- | --- | --- | --- |
| **Sensorimotor** | Y - OA HF | 0.033 | 0.036 | 0.932 | 0.354 |
|  | Y - OA LF | 0.128 | 0.032 | 4.036 | < 0.001 |
|  | OA LF - OA- HF | 0.095 | 0.030 | 3.218 | 0.004 |
| **Vestibular** | Y - OA HF | 0.080 | 0.020 | 4.039 | < 0.001 |
|  | Y - OA LF | 0.116 | 0.018 | 6.632 | < 0.001 |
|  | OA LF - OA- HF | 0.036 | 0.016 | 2.236 | 0.028 |
| **DLPFC** | Y - OA HF | 0.064 | 0.014 | 4.632 | < 0.001 |
|  | Y - OA LF | 0.070 | 0.012 | 5.718 | < 0.001 |
|  | OA LF - OA- HF | 0.006 | 0.011 | 0.527 | 0.600 |
| **ACC** | Y - OA HF | 0.021 | 0.007 | 3.108 | 0.005 |
|  | Y - OA LF | 0.037 | 0.006 | 6.200 | < 0.001 |
|  | OA LF - OA- HF | 0.016 | 0.006 | 2.903 | 0.005 |
| **DMN** | Y - OA HF | 0.035 | 0.014 | 2.406 | 0.036 |
|  | Y - OA LF | 0.060 | 0.013 | 4.711 | < 0.001 |
|  | OA LF - OA- HF | 0.026 | 0.012 | 2.152 | 0.036 |
| **Visual** | Y - OA HF | 0.094 | 0.023 | 4.131 | < 0.001 |
|  | Y - OA LF | 0.089 | 0.020 | 4.412 | < 0.001 |
|  | OA LF - OA- HF | -0.005 | 0.019 | -0.272 | 0.786 |

**Table 4.** Statistical output for post-hoc pairwise group comparisons in between-network connectivity scores. P-values are adjusted for multiple comparisons using the Holm-Bonferroni method. Y = Young adults. OA-HF = Older adults with high physical function. OA-LF = Older adults with low physical function. DLPFC = Dorsolateral Prefrontal Cortex. ACC = Anterior Cingulate Cortex. DMN = Default Mode Network.

| Network | Contrast | Estimate | SE | t-ratio | p-value |
| --- | --- | --- | --- | --- | --- |
| **Sensorimotor** | Y - OA HF | 0.012 | 0.008 | 1.488 | 0.140 |
|  | Y - OA LF | 0.026 | 0.007 | 3.697 | 0.001 |
|  | OA LF - OA- HF | 0.014 | 0.006 | 2.175 | 0.064 |
| **Vestibular** | Y - OA HF | 0.013 | 0.008 | 1.649 | 0.102 |
|  | Y - OA LF | 0.026 | 0.007 | 3.936 | 0.001 |
|  | OA LF - OA- HF | 0.014 | 0.006 | 2.237 | 0.055 |
| **DLPFC** | Y - OA HF | 0.008 | 0.006 | 1.440 | 0.153 |
|  | Y - OA LF | 0.017 | 0.005 | 3.393 | 0.003 |
|  | OA LF - OA- HF | 0.009 | 0.005 | 1.907 | 0.119 |
| **ACC** | Y - OA HF | 0.008 | 0.005 | 1.485 | 0.141 |
|  | Y - OA LF | 0.015 | 0.004 | 3.461 | 0.002 |
|  | OA LF - OA- HF | 0.008 | 0.004 | 1.924 | 0.114 |
| **DMN** | Y - OA HF | 0.011 | 0.006 | 1.996 | 0.097 |
|  | Y - OA LF | 0.020 | 0.005 | 3.922 | 0.001 |
|  | OA LF - OA- HF | 0.008 | 0.005 | 1.800 | 0.097 |
| **Visual** | Y - OA HF | 0.010 | 0.006 | 1.588 | 0.132 |
|  | Y - OA LF | 0.019 | 0.006 | 3.518 | 0.002 |
|  | OA LF - OA- HF | 0.010 | 0.005 | 1.861 | 0.132 |

Group x Laterality interaction effect on within-network connectivity was significant for the vestibular network, but not for sensorimotor, dorsolateral prefrontal cortex, and anterior cingulate cortex networks (Interaction effect, Vestibular: F(2,98) = 9.39, p<0.001; Interaction effect, Sensorimotor: F(2,98) = 0.22, p = 0.807; Interaction effect, Dorsolateral prefrontal cortex: F(2,98) = 0.31, p = 0.733; Interaction effect, Anterior cingulate cortex: F(2,98) = 1.00, p = 0.373). Post-hoc tests on within-group differences in left and right vestibular within-network connectivity showed that all groups had lower connectivity in the left network compared to the right (Supplementary table 2; all p<0.050); Left and right within-network connectivity differences were largest in the younger adult group for the vestibular network (Young adults group estimate = –0.050; Older adults with higher physical function group estimate = –0.014; Older adults with lower physical function group estimate= –0.017).

Default mode within-network connectivity was different between all groups (F(2, 98) = 11.34, p < 0.001; Figure 3 C-D); younger adults had higher within-network connectivity compared to both older adult groups, and older adults with higher physical function had higher within-network connectivity compared to older adults with lower physical function. Visual within-network connectivity was higher in younger adults compared to both older adults groups, but group differences between older adults with higher and lower physical function was not statistically significant (Group effect: F(2,98) = 11.27, p < 0.001).

In general, between-network connectivity was higher in younger adults compared to older adults. Sensorimotor, vestibular, dorsolateral prefrontal cortex, and anterior cingulate cortex between-network connectivity were lower in older adults with lower physical function compared to young adults, but between-network connectivity was not significantly different between younger adults and older adults with higher physical function, and between older adult sub-groups (Group effect, Sensorimotor: F(2,98) = 7.40, p = 0.001; Group effect, Vestibular: F(2,98) = 8.30, p < 0.001; Group effect, Dorsolateral prefrontal cortex: F(2,98) = 6.15, p =0.003; Group effect, Anterior cingulate cortex: F(2,98) = 6.37, p =0.002). Between-network segregation was higher in the left compared to the right for sensorimotor, vestibular, dorsolateral prefrontal cortex, and anterior cingulate cortex networks (Sensorimotor: F(1,98) = 268.77, p < 0.001; Vestibular = F(1,98) = 275.88, p < 0.001; Dorsolateral prefrontal cortex: F(1,98) = 74.48, p < 0.001; Anterior cingulate cortex: F(1,98) = 14.59, p < 0.001). Group x Laterality interaction effect on between-network connectivity was significant for the sensorimotor, vestibular, and dorsolateral prefrontal cortex networks, but not for the anterior cingulate cortex network (Interaction effect, Sensorimotor: F(2,98) = 5.40, p = 0.006; Interaction effect, Vestibular: F(2,98) = 6.61, p = 0.002; Interaction effect, Dorsolateral prefrontal cortex: F(2,98) = 4.25, p = 0.017; Interaction effect, Anterior cingulate cortex: F(2,98) = 1.00, p = 0.373). Post-hoc tests on between-network connectivity showed that all groups had higher between-network connectivity in the left network compared to the right (Supplementary table 3; all p’s ≦ 0.001); Left and right between-network connectivity differences were largest in the younger adult group (Sensorimotor: Young adults group estimate = 0.026; Older adults with higher physical function group estimate = 0.024; Older adults with lower physical function group estimate = 0.016; Vestibular: Younger adults group estimate = 0.017; Older adults with higher physical function group estimate = 0.016; Older adults with lower physical function estimate = 0.010; Dorsolateral prefrontal cortex: Younger adults group estimate = 0.002; Older adults with higher physical function group estimate = 0.002; Older adults with lower physical function group estimate = 0.001).

Default mode and visual between-network connectivity was lower in older adults with lower physical function compared to younger adults, but not significantly different between younger adults and older adults with higher physical function, and between older adult sub-groups (Default mode: F(2,98)= 7.87, p = 0.001; Visual: F(2,98) = 6.50, p = 0.002; Figure 3 E-F).

### 3.3. Association with walking speed and working memory performance: Age groups

Canonical correlation analysis with age groups showed that only the first canonical correlation was statistically significant (r = 0.627, p < 0.001) as shown in Table 5. Figure 5 presents the standardized canonical coefficients for the first dimension. In the Function variable dataset the canonical correlation was most strongly influenced by age groups (Coef. = –0.761), walking speed (Coef. = –0.567), and working memory (Coef. = 0.193). For network segregation, the canonical correlation was most strongly influenced by right vestibular (Coef. = –0.434), left sensorimotor (Coef. = –0.384), default mode network (Coef. = 0.270), and right dorsolateral prefrontal cortex (Coef. = –0.199). This observation demonstrates that younger adults with faster gait speed had higher network segregation in right vestibular, left sensorimotor, and right dorsolateral prefrontal cortex networks, and lower network segregation in the default mode network.

**Figure 5.**
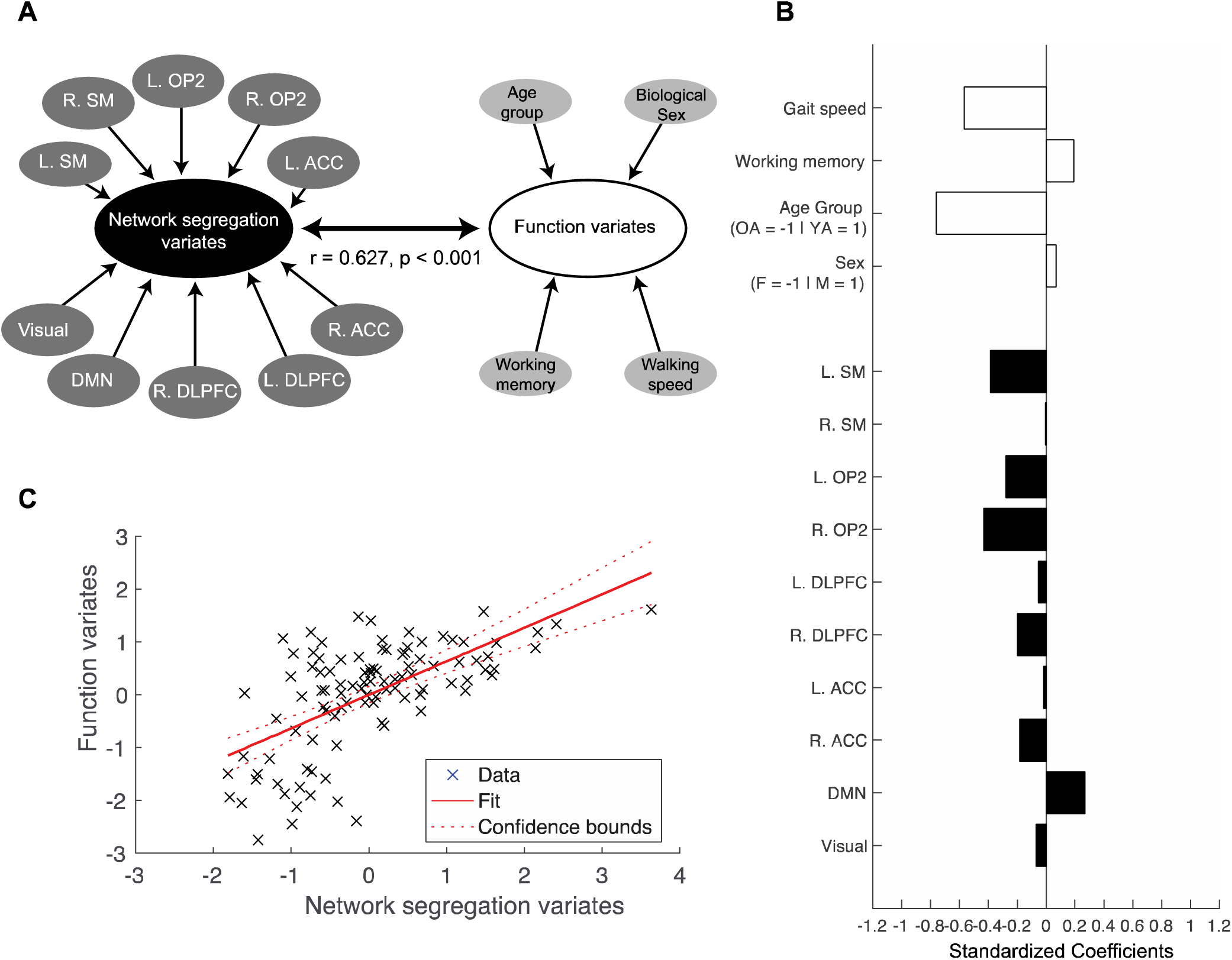
Canonical correlation analysis for age groups. A. Canonical correlation analysis (CCA) was conducted to assess the relationship between the ‘function dataset’, which included walking speed, cognitive function, age group, and biological sex, and the ‘network segregation dataset’, which included network segregation scores from the 10 identified networks. First dimension showed significant association of variance between the two datasets. B. Standardized correlation coefficient for the first dimension. White bars = Coefficients for the Function dataset; Black bars = Coefficients for the Network segregation dataset. C. Canonical variates for the first dimension. Canonical variates are values of the canonical variables which is computed by multiplying the normalized variables by the canonical coefficients. L = Left; R = Right; SM = Sensorimotor; OP2 = Vestibular; DLPFC = Dorsolateral prefrontal cortex; ACC = Anterior cingulate cortex; DMN = Default mode network.

**Table 5.** Statistical output for canonical correlation analysis with age groups. Dimension reduction results from Wilks lambda are presented.

| Dimension | Corr. | Wilks | F | df1 | df2 | p |
| --- | --- | --- | --- | --- | --- | --- |
| 1-4 | 0.627 | 0.408 | 2.24 | 40 | 335.54 | < 0.001 |
| 2-4 | 0.425 | 0.672 | 1.41 | 27 | 260.57 | 0.093 |
| 3-4 | 0.376 | 0.821 | 1.17 | 16 | 180.00 | 0.297 |
| 4-4 | 0.211 | 0.955 | 0.61 | 7 | 91.00 | 0.749 |

We tested two multiple linear regression models with age groups: (1) Walking speed and (2) working memory, both with biological sex covaried out. For walking speed, univariate linear regressions determined the left and right sensorimotor, left and right vestibular, left and right dorsolateral prefrontal cortex, left and right anterior cingulate cortex, default mode, and visual network segregation, and left anterior cingulate cortex network segregation x Age group interaction to be considered in the step-wise multiple regression model (Supplementary figure 1). Multiple linear regression with walking speed was statistically significant (F(6,93) = 8.71, p < 0.001, r^2^ = 360). Left sensorimotor (p = 0.003) and left anterior cingulate cortex network segregation (p = 0.030), and interaction of left anterior cingulate cortex network segregation x Age group (p = 0.002) were significant predictors. Age group (p = 0.286), left vestibular (p = 0.148), and right anterior cingulate cortex network segregation (p = 0.114) were not significant predictors, but contributed to the model (Table 6).

**Table 6.** Statistical output for step-wise multiple regression with walking speed (controlled for biological sex) in young and older adults (F(6, 93) = 8.71, p < 0.001, r^2^ = 0.36). Analysis included 20 young adults and 80 older adults (1 young adult and 1 older adult with higher physical function excluded due to missing walking speed data). Y = Young adults. ACC = Anterior Cingulate Cortex.

|  | Estimate | Standardized | SE | t-value | p-value |
| --- | --- | --- | --- | --- | --- |
| **Intercept** | -0.28 | -- | 0.06 | -4.89 | < 0.001 |
| **Left Sensorimotor** | 0.40 | 0.34 | 0.11 | 3.75 | < 0.001 |
| **Left Vestibular** | 0.20 | 0.14 | 0.14 | 1.46 | 0.148 |
| **Left ACC** | -0.29 | -0.27 | 0.13 | -2.21 | 0.030 |
| **Right ACC** | 0.20 | 0.19 | 0.12 | 1.59 | 0.114 |
| **Age group (Y)** | -0.08 | -0.17 | 0.07 | -1.08 | 0.283 |
| **Left ACC x Age group (Y)** | 0.78 | 0.50 | 0.24 | 3.19 | 0.002 |

For working memory, univariate linear regressions determined left and right sensorimotor, left and right vestibular, left and right dorsolateral prefrontal cortex, left and right anterior cingulate cortex, and right anterior cingulate cortex x age group interaction to be considered in the step-wise multiple regression model (Supplementary figure 2). Multiple linear regression with working memory was statistically significant (F(3,72) = 11.70, p < 0.001, r^2^ = 0.328); however only left sensorimotor network segregation (p = 0.428), age group (p = 0.272) and the interaction of left sensorimotor network segregation x age group (p = 0.045) were contributors to the model (i.e., the model was a univariate model) (Table 7).

**Table 7.** Statistical output for step-wise multiple regression with working memory (controlled for biological sex) in young and older adults (F(3, 72) = 11.70, p < 0.001, r^2^ = 0.33). Analysis included 18 young adults and 58 older adults (3 young adults and 23 older adults excluded due to missing working memory scores). Y = Young adults.

|  | Estimate | Standardized | SE | t-value | p-value |
| --- | --- | --- | --- | --- | --- |
| **Intercept** | -0.17 | -- | 1.05 | -0.17 | 0.869 |
| **Left Sensorimotor** | -1.69 | -0.08 | 2.12 | -0.80 | 0.428 |
| **Age group (Y)** | -5.00 | -0.66 | 4.52 | -1.11 | 0.272 |
| **Left Sensorimotor x Age group (Y)** | 16.46 | 1.24 | 8.06 | 2.04 | 0.045 |

### 3.4. Association with walking speed and working memory: Physical function groups

Canonical correlation analysis with physical function groups showed that the first (r = 0.627, p = 0.002) was significant. Dimension reduction showed that the second, third and fourth dimensions did not significantly explain shared variance (Table 8). Figure 6 presents the standardized canonical coefficients for the first dimension. For the first canonical dimension, canonical correlation was most strongly influenced by walking speed (Coef. = –0.698), working memory performance (Coef. = 0.451), and physical function groups (Coef. = –0.396) from the function dataset. From the network segregation dataset, left sensorimotor (Coef. = –0.619), default mode (Coef. = –0.413), left dorsolateral prefrontal cortex (Coef. = 0.381), visual networks (Coef. = 0.347) most strongly influenced the canonical correlation. This observation demonstrates that older adults in the lower physical function group with slower walking speed and higher working memory had lower network segregation in the left sensorimotor and default mode networks and higher segregation in the left dorsolateral prefrontal cortex and visual networks.

**Figure 6.**
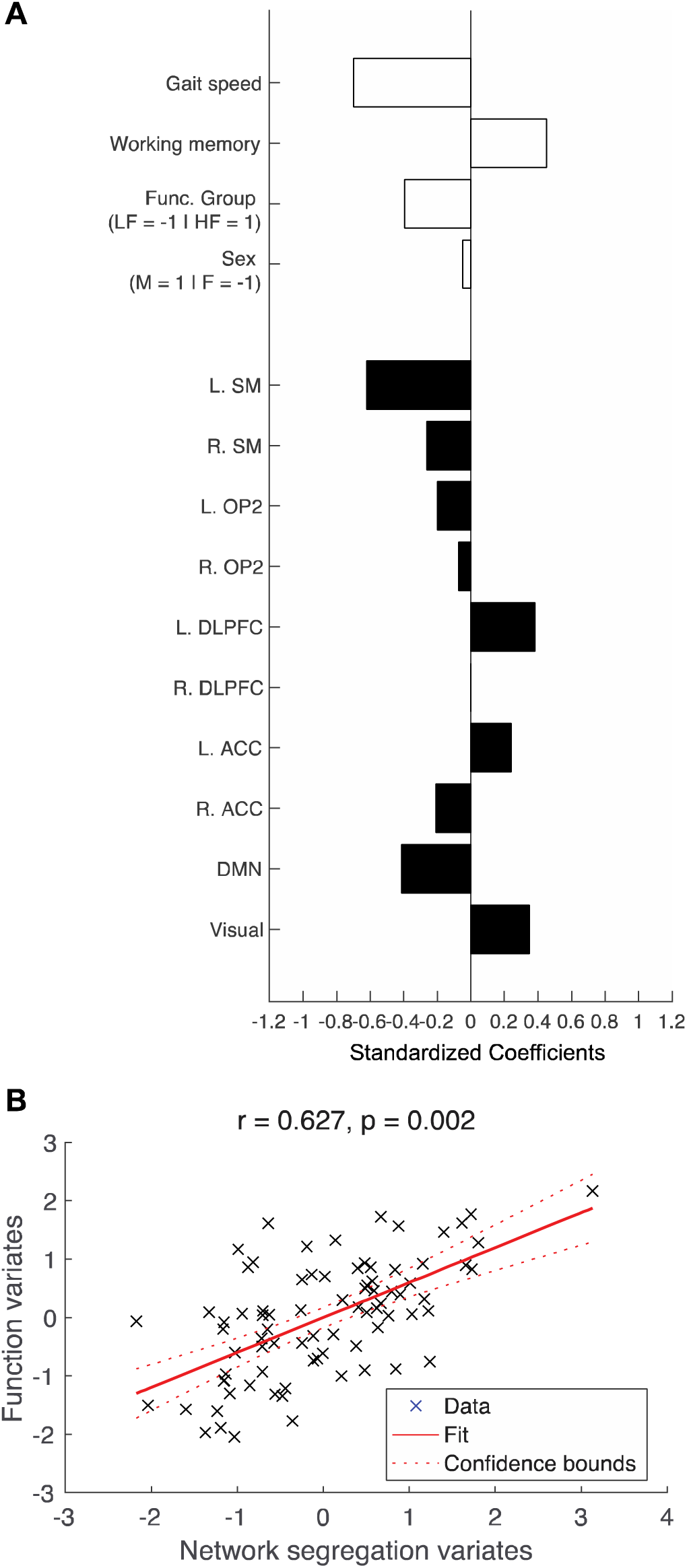
Canonical correlation analysis for physical function groups. Canonical correlation analysis (CCA) was conducted to assess the relationship between the ‘function dataset’, which included walking speed, cognitive function, physical function group, and biological sex, and the ‘network segregation dataset’, which included network segregation scores from the 10 identified networks. First dimension showed significant association of variance between the two datasets. A. Standardized correlation coefficient for the first and second dimensions. White bars = Coefficients for the Function dataset; Black bars = Coefficients for the Network segregation dataset. B. Canonical variates for the first dimension. Canonical variates are values of the canonical variables which is computed by multiplying the normalized variables by the canonical coefficients. L = Left; R = Right; SM = Sensorimotor; OP2 = Vestibular; DLPFC = Dorsolateral prefrontal cortex; ACC = Anterior cingulate cortex; DMN = Default mode network.

**Table 8.** Statistical output for canonical correlation analysis with physical function groups (older adult participants only). Dimension reduction results from Wilks lambda are presented.

| Dimension | Corr. | Wilks | F | df1 | df2 | p |
| --- | --- | --- | --- | --- | --- | --- |
| 1-4 | 0.627 | 0.374 | 1.90 | 40 | 255.91 | 0.002 |
| 2-4 | 0.472 | 0.615 | 1.34 | 27 | 199.24 | 0.135 |
| 3-4 | 0.425 | 0.792 | 1.07 | 16 | 138.00 | 0.391 |
| 4-4 | 0.185 | 0.966 | 0.35 | 7 | 70.00 | 0.926 |

For walking speed with older adults only, univariate linear regressions determined the left and right sensorimotor, left and right vestibular, left and right dorsolateral prefrontal cortex, left and right anterior cingulate cortex, default mode, and visual network segregation to be considered in the step-wise multiple regression model (Supplementary figure 3). Post-hoc multiple linear regression analysis with walking function was statistically significant (F(5,74) = 9.56, p < 0.001, r^2^ = 0.392). Right sensorimotor (p = 0.009), left anterior cingulate cortex (p = 0.012), and right anterior cingulate cortex network segregation (p = 0.036), and functional group (p < 0.001) were significant predictors. Right vestibular network segregation (p = 0.108) was not a significant predictor but it contributed to the model. The correlation coefficient estimate for the left anterior cingulate cortex was negative, while all other correlation coefficients were positive (Table 9). For working memory, none of the network segregation scores were significantly associated in the univariate analysis, so no multiple linear regression model was tested (Supplementary figure 4).

**Table 9.** Statistical output for step-wise multiple regression with walking speed (controlled for biological sex) in older adults with high and low physical function (younger adults were excluded from this analysis) (F(5, 74) = 9.56, p < 0.001, r^2^ = 0.39). Analysis included 26 older adults with high physical function and 54 older adults with low physical function (1 older adult with higher physical function excluded due to missing walking speed data). HF = Older adults with high physical function. ACC = Anterior Cingulate Cortex.

|  | Estimate | Standardized | SE | t-value | p-value |
| --- | --- | --- | --- | --- | --- |
| **Intercept** | -0.32 | -- | 0.07 | -4.74 | < 0.001 |
| **Right Sensorimotor** | 0.25 | 0.26 | 0.09 | 2.67 | 0.009 |
| **Right Vestibular** | 0.21 | 0.16 | 0.13 | 1.63 | 0.108 |
| **Left ACC** | -0.32 | -0.31 | 0.12 | -2.58 | 0.012 |
| **Right ACC** | 0.26 | 0.26 | 0.12 | 2.14 | 0.036 |
| **Functional group (HF)** | 0.14 | 0.38 | 0.03 | 3.98 | < 0.001 |

## 4. DISCUSSION

We found that network segregation in sensorimotor, vestibular, dorsolateral prefrontal cortex, anterior cingulate cortex, and visual networks were higher in younger adults than older adults, but only sensorimotor, and vestibular network segregation was different between older adults with high and low physical function (Aim 1a). We observed laterality differences in segregation scores for some networks, but no group by laterality interactions (Aim 1b). We observed multivariate associations between working memory and walking speed with network segregation scores (Aim 2). For young and old age groups, canonical correlation analysis demonstrated that right vestibular, left sensorimotor, default mode network, and right dorsolateral predeontal network segregation largely influenced the variability in walking speed observed. Working memory was associated with interaction of left sensorimotor cortex network segregation and age groups. For older adults with high and low physical function, canonical correlation analysis demonstrated that higher walking speed were largely influenced by higher segregation in the left sensorimotor and default mode networks, and lower segregation in the left dorsolateral prefrontal cortex, and visual networks.

### 4.1. Network segregation in mobility-related networks is different between older adults with high and low physical function

We observed that older adults with higher physical function had significantly higher segregation in the sensorimotor and vestibular networks than their peers with lower physical function (as determined with the short physical performance battery score). Importantly, while aging studies have consistently demonstrated that older adults have decreased network segregation compared to younger adults (Cassady et al., 2019; Chan et al., 2014; Damoiseaux, 2017; King et al., 2018), we observed that sensorimotor, anterior cingulate, and default mode network segregation was not significantly different between younger adults and older adults with high physical function (and dorsolateral prefrontal cortex with borderline significance between younger adults and older adults with higher physical function: p = 0.49, Table 2). Our results demonstrated that older adults with higher physical function had fewer differences with younger adults compared to older adults with lower physical function and that mobility-related networks (sensorimotor and vestibular) were more segregated in higher functioning older adults compared to the lower functioning older adults. Older adults with higher physical function may have more resilience to decrease in network segregation due results from previous studies suggesting that they have better cardiovascular function and cerebrovascular health (Smith et al., 2021). Or, similar network segregation in older adults to younger adults may help intrinsic brain activity (e.g., by more efficient communication of the brain, leading to better functioning motor-cognitive integration) leading to higher physical function. Whether the network segregation differences between the younger adults and older adult sub-groups reflect cause or effect of physical function differences is unknown and outside the scope of this study. However, network segregation appears to be a key correlate of physical function in older adults.

In line with our hypothesis and previous studies (Cassady et al., 2020; Cassady et al., 2019), sensorimotor network segregation was higher in younger adults compared to older adults. When we examined several different networks, networks that are important for mobility function such as sensorimotor, vestibular, and visual networks demonstrated group differences in network segregation, but networks that are typically involved in more cognitive functions-the dorsolateral prefrontal cortex, anterior cingulate cortex, also demonstrated differences between groups. Our results suggest that mobility-related networks (i.e., sensorimotor and vestibular networks which showed strong differences between younger adults and older adults with lower physical function) may be more susceptible to age-related changes in network segregation compared to cognitive function networks. This is in line with behavioral studies that showed that mobility declines precede cognitive decline in older adults and neurodegenerative disease such as Alzheimer’s and mild cognitive impairment (Buracchio et al., 2010; Merchant et al., 2021; Skillbäck et al., 2022). Moreover, recent studies on age differences in brain structure reveal the largest age effects in the pre and post central gyri, the thalamus, striatum, and cerebellum (Hupfeld et al., 2022; Taubert et al., 2020).

We expected within-network connectivity to be higher and between-network connectivity to be lower in older adults compared to younger adults. However, in general, younger adults had higher connectivity in both within-network and between-network connectivity. This suggests that the lower network segregation scores in older adults are driven more by within-than between-network connectivity differences. Interestingly, between-network connectivity was not significantly different in any of the networks between young adults and older adults with higher physical function. This may suggest that between-network connectivity is less susceptible to age-related brain connectivity differences.

### 4.2. Laterality in network segregation is similar between younger and older adults

We showed group x laterality interactions in both the within-network connectivity and between network connectivity. Left and right connectivity differences were largest in the younger adults (as shown by the post-hoc estimates) for vestibular network within-network connectivity, and sensorimotor, vestibular, and dorsolateral prefrontal cortex between-network connectivity. The right vestibular network (which is the dominant side network) had stronger within-network connectivity compared to the left side. This is in line with expectation and previous work in task-based functional MRI showing reduced lateralization in older adults. Reduced lateralization of the cortex has been shown in seminal work in cognitive processing, resulting in the hemispheric asymmetry reduction in older adults (HAROLD) conceptual model (Cabeza, 2002; Cabeza et al., 2002). However, the group x laterality interactions were not present in the segregation scores. This may suggest that the between-network connectivity may be compensating for the changes in within-network connectivity.

It may also be that decreased lateralization with increased age is primarily evident in task-based functional MRI data rather than resting state network segregation scores. The differences in age-related laterality outcomes between task-based and resting-state functional MRI highlight the complexity of brain laterality. Task-based fMRI demonstrates strong lateralization effects due to specific cognitive demands brain during different tasks (Cabeza, 2002; Cabeza et al., 2002). In contrast, resting-state fMRI characterizes the brain’s intrinsic activity without any external stimuli, leading to diffuse communication patterns in the brain. Furthermore, networks are defined by nodes that are functionally connected and often span both hemispheres, leading to increased complexity in interpreting ‘laterality’ of functional networks. Our study showing lack of interaction effects in age groups and lateralization suggests that the variability in laterality observed in resting-state data may differ significantly from task-induced activation patterns.

### 4.3. Resting state network segregation may have differential roles in function

In general, for both of our canonical correlation models, majority of the network segregation was positively associated with walking speed. This is in support of previous brain-behavior relationships that showed sensorimotor network segregation positively correlated with general sensorimotor function in both younger and older adults (Cassady et al., 2020; Cassady et al., 2019), and higher network segregation is predictive of greater intervention-related cognitive gains (Cohen and D’Esposito, 2016; Gallen et al., 2016; Gallen and D’Esposito, 2019).

However, our canonical correlation analysis demonstrated that not all resting state networks are positively correlated to walking speed and working memory. We expected network segregation to be positively correlated to behavioral function as studies have consistently demonstrated lower network segregation in older adults compared to younger adults, suggestive of decreased efficiency of functional networks (Deery et al., 2023). Studies of cognitive aging have supported the scaffolding theory of aging and cognition (STAC), which suggests that the brain adapts and reorganizes to recruit additional neural resources to preserve cognitive function, which is a result of functional and structural challenges that occur with aging (Park and Reuter-Lorenz, 2009; Reuter-Lorenz and Park, 2014). The networks with the opposite canonical correlation coefficients compared to working memory and walking speed in our model support that STAC may also apply to our multi-function model including walking and cognition.

In our multiple regression model with walking speed, there was a significant age group x left anterior cingulate cortex network segregation interaction, in which the correlation coefficient was positive in younger adults, while the correlation coefficient was negative in older adults (Also shown in Supplementary figure 1G). This is in contrast to our initial hypothesis and provides novel insight that not all lower segregation is related to decreased function. Specifically, the left anterior cingulate cortex network segregation may be playing a compensatory role in older adults with regards to walking function. The left anterior cingulate cortex plays crucial roles in cognition, emotional processing, and error monitoring (Gwin et al., 2011; Peterson and Ferris, 2019; Sipp et al., 2013). The negative correlation of left anterior cingulate cortex may demonstrate that older adults recruit additional neural resources for the left anterior cingulate cortex network for diffuse connectivity to compensate for the lower network segregation in left sensorimotor, left vestibular, and right anterior cingulate cortex networks underlying slower walking function.

We expected networks with seed regions of primarily cognitive roles (i.e., dorsolateral prefrontal cortex and anterior cingulate cortex) to be associated with working memory. However, our results demonstrate that in young and older adults with varying physical function the interaction of sensorimotor network segregation and age groups was associated with working memory. In older adults, higher left sensorimotor network segregation was associated with lower working memory, while in younger adults, higher left sensorimotor network segregation was associated with higher working memory. This may suggest that in younger adults, higher left network segregation indicates a ‘healthier’ brain with higher working memory. However, in older adults, higher left sensorimotor network was associated to lower working memory, which may suggest that the left sensorimotor network segregation is compensating for other functions in the brain such as functional activation in the dorsolateral prefrontal cortex.

Our results demonstrating association with network segregation and behavior may suggest potential interventions to modulate resting state network segregation to increase functional performance in walking or working memory. Recently, transcranial direct stimulation has been shown to increase resting state network segregation (Iordan et al., 2022). Clinical trials to modulate resting state network segregation have mainly been with cognitively impaired populations (i.e., Alzheimer’s disease and related dementias and mild cognitive impairment) as this population has reduced network segregation (Ewers et al., 2021; Fu et al., 2022; Iordan et al., 2022; Steward et al., 2023). These outcomes are often short-lived (Aksu et al., 2024) and results on the efficacy of transcranial direct current stimulation are mixed in the literature (Duffy et al., 2024). However, our results demonstrating association with walking speed and resting state network segregation serves as an initial step to suggest that network segregation may serve as a potential therapeutic target to maintain or increase walking speed in typical older adults. The potential causal role of network segregation for use in therapeutic interventions remains speculative within our current cross-sectional study.

Nevertheless, future longitudinal studies looking at network segregation and behavior associations in aging remains crucial for further understanding the potential implication of network segregation for therapeutic outcomes to maintaining and enhancing physical function in older adults.

### 4.4. Limitations

As mentioned in the methods section, this study was part of a larger NIH U01 study at the University of Florida (U01AG061389) (Clark et al., 2019). Due to how the larger study was designed, the brain scans and behavioral tests took place on different days (Average number of days between brain scans and behavioral tests: 81.5 ± 94.8 days). In addition, data collection for the study was interrupted due to the COVID-19 pandemic. This may be a limitation in our current study because there is a variability in the time between the brain scan collection date and the behavioral tests collection date. However, it should be noted that resting-state brain network segregation is generally stable overtime / reliable as long as sufficient clean data is used (Pierce et al., 2024), even despite aging declines (Rosenberg et al., 2020). In addition, as outlined in our protocol paper (Clark et al., 2019), we had strict exclusion/inclusion criteria, and all participants were required to be free from any major neurological injuries. Therefore, we expect little to no variability in resting state brain network segregation due to the time between the brain scan collection date and the behavioral tests.

Another limitation of the study is that the older adult cohort who participated in this study represents a relatively high functioning population. Even those in the low functioning group who scored 10 or less on the short performance battery score were required to be able to walk 400 m without sitting or assistance of a person or a walking device as part of the inclusion criteria. Typical walking speeds for community-dwelling older adults range from 0.9-1.30 m/s(Bohannon et al., 1996; Krishnamurthy and Verghese, 2006), and range from 0.23-0.50 m/s in older adults in hospital or rehabilitation settings (Friedman et al., 1988; Purser et al., 2005). The overall higher physical function of our older adults may limit the generalizability of our findings to a broader population. However, our mean gait speed is in the range of older adults living in a community setting, and the group differences between high and low functioning groups that we observed in this study emphasize the potential implications of physical health on resting-state brain connectivity (even within community-dwelling adults). A challenge of interpreting studies in aging is the wide variability in function of older adults. To facilitate interpretation of results, we included a table on participant characteristics. Our study offers a valuable initial insight into network segregation in aging. Future research with more diverse samples, including older adults with lower physical function, remains crucial to fully understand the scope and implications of our findings. Readers should take caution in generalizing our findings to a broader population for these reasons.

### 4.5. Conclusion

Resting state brain network segregation is emerging as a metric that is sensitive to age, mobility and cognitive function, and is predictive of cognitive gains from various interventions (Cohen and D’Esposito, 2016; Gallen et al., 2016; Gallen and D’Esposito, 2019). A greater understanding of this metric and its behavioral correlates may help identify biomarkers related to function. In this study, we demonstrated age and physical function group differences in network segregation and laterality in network segregation, and we identified a multivariate brain-behavior model that showed differential roles of network segregation specific to function. This multivariate model is novel and significant because we show that predominantly higher network segregation is associated with higher walking speed and working memory. These findings highlight the importance of resting state brain network segregation to walking and working memory function in older and younger adults.

## Data availability

Data will be made available upon request. Coordinates for resting state network segregation analysis are provided in Supplementary table 1.

