## Supplementary material for "Resting state brain network segregation is associated with walking speed and working memory in older adults"

Supplementary table 1. MNI coordinates for the nodes used in this study. OP2 = Vestibular; DLPFC = Dorsolateral prefrontal cortex; ACC = Anterior cingulate cortex.

Supplementary table 2. Statistical output for post-hoc within-group, between-conditions pairwise comparisons between left and right vestibular within-network connectivity. P-values are adjusted for multiple comparisons using the Holm-Bonferroni method. YA = Young adults. OA-HF = Older adults with high physical function. OA-LF = Older adults with low physical function.

|  |  |  |  |  | 95% confidence interval | |
| --- | --- | --- | --- | --- | --- | --- |
|  |  | p-value | t-statistic | estimate | Low | High |
| Vestibular within-network connectivity | YA | < 0.001 | -5.06 | -0.050 | -0.070 | -0.029 |
|  | OA- HF | 0.014 | -2.65 | -0.014 | -0.025 | -0.003 |
|  | OA- LF | < 0.001 | -4.46 | -0.017 | -0.025 | -0.009 |

Supplementary table 3. Statistical output for post-hoc within-group, between-conditions pairwise comparisons between left and right sensorimotor, vestibular, and dorsolateral prefrontal cortex (DLPFC) between-network connectivity. P-values are adjusted for multiple comparisons using the Holm-Bonferroni method. YA = Young adults; OA-HF = Older adults with high physical function; OA-LF = Older adults with low physical function; DLPFC = Dorsolateral prefrontal cortex.

|  |  |  |  |  | 95% confidence interval | |
| --- | --- | --- | --- | --- | --- | --- |
|  |  | p-value | t-statistic | estimate | Low | High |
| Sensorimotor between-network connectivity | YA | < 0.001 | 7.55 | 0.026 | 0.019 | 0.033 |
|  | OA- HF | < 0.001 | 7.90 | 0.024 | 0.018 | 0.030 |
|  | OA- LF | < 0.001 | 12.80 | 0.016 | 0.014 | 0.019 |
| Vestibular between-network connectivity | YA | < 0.001 | 8.03 | 0.017 | 0.013 | 0.021 |
|  | OA- HF | < 0.001 | 7.96 | 0.016 | 0.012 | 0.020 |
|  | OA- LF | < 0.001 | 12.20 | 0.010 | 0.008 | 0.012 |
| DLPFC between-network connectivity | YA | 0.001 | 4.13 | 0.002 | 0.001 | 0.003 |
|  | OA- HF | < 0.001 | 5.16 | 0.002 | 0.001 | 0.002 |
|  | OA- LF | < 0.001 | 4.35 | 0.001 | 0.000 | 0.001 |

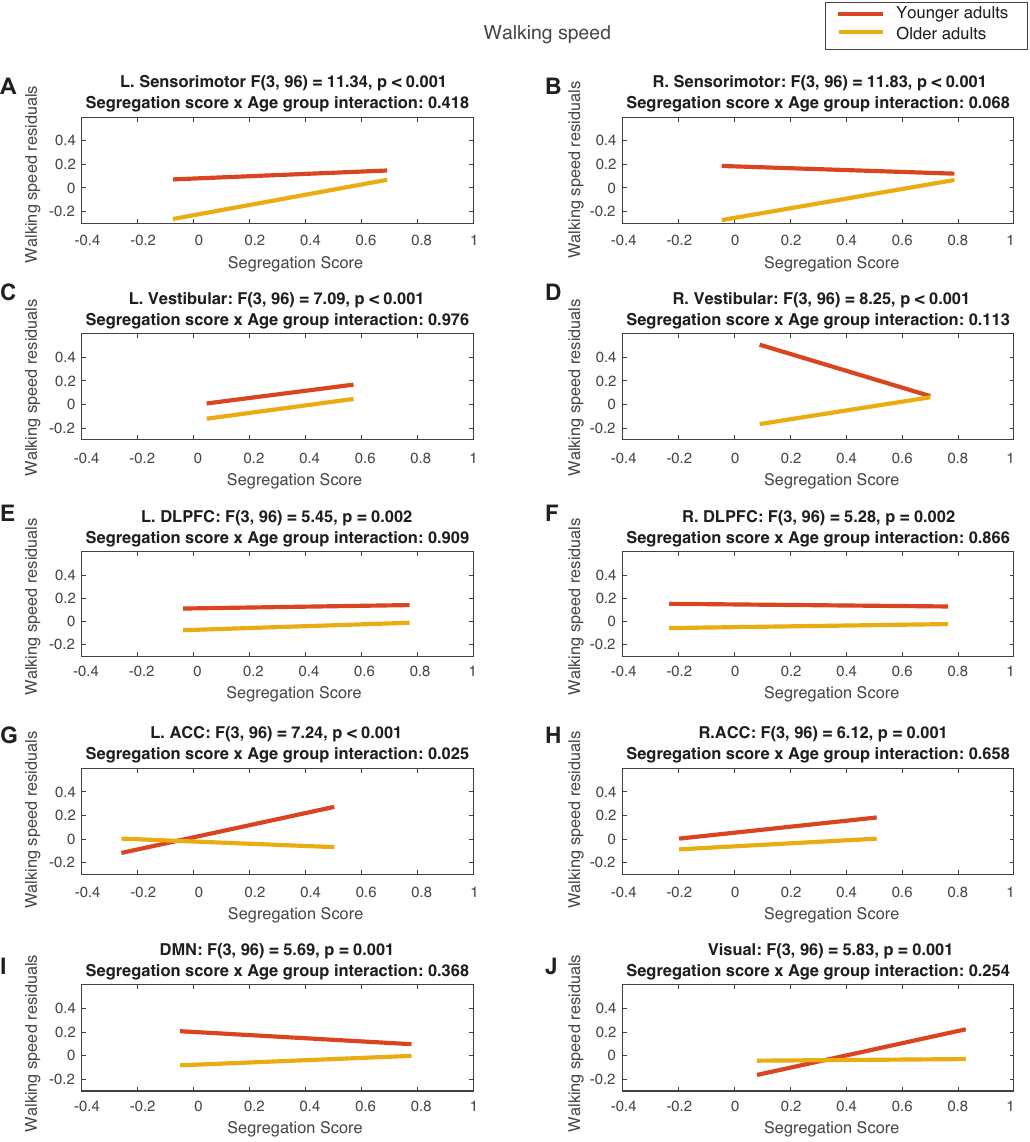

Supplementary figure 1. Univariate linear regression between walking speed corrected for biological sex and segregation scores for each network, with age groups. Lines represent best-fit line. DLPFC = Dorsolateral prefrontal cortex; ACC = Anterior cingulate cortex; DMN = Default mode network.

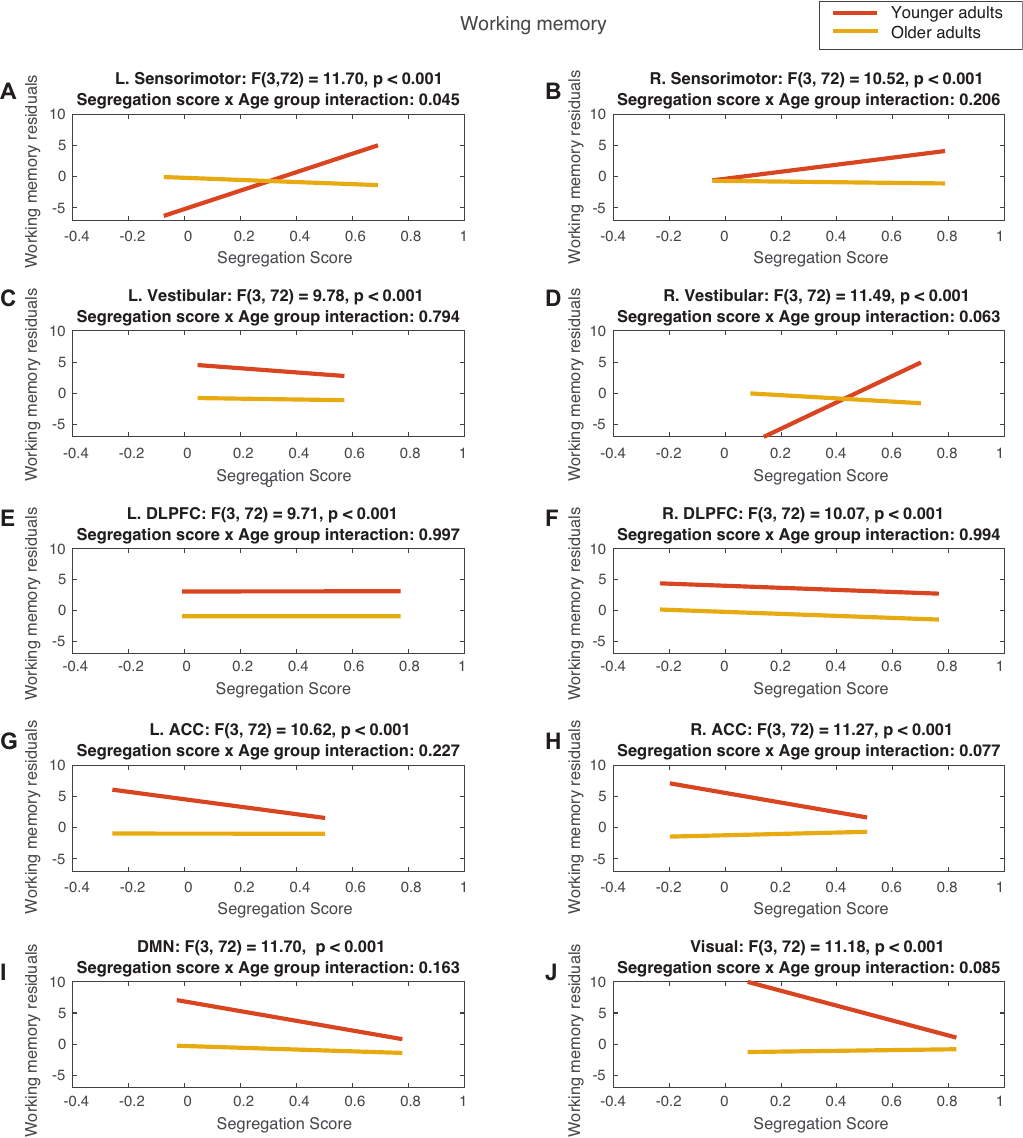

Supplementary figure 2. Univariate linear regression between working memory corrected for biological sex and segregation scores for each network, with age groups. Lines represent best-fit line. DLPFC = Dorsolateral prefrontal cortex; ACC = Anterior cingulate cortex; DMN = Default mode network.

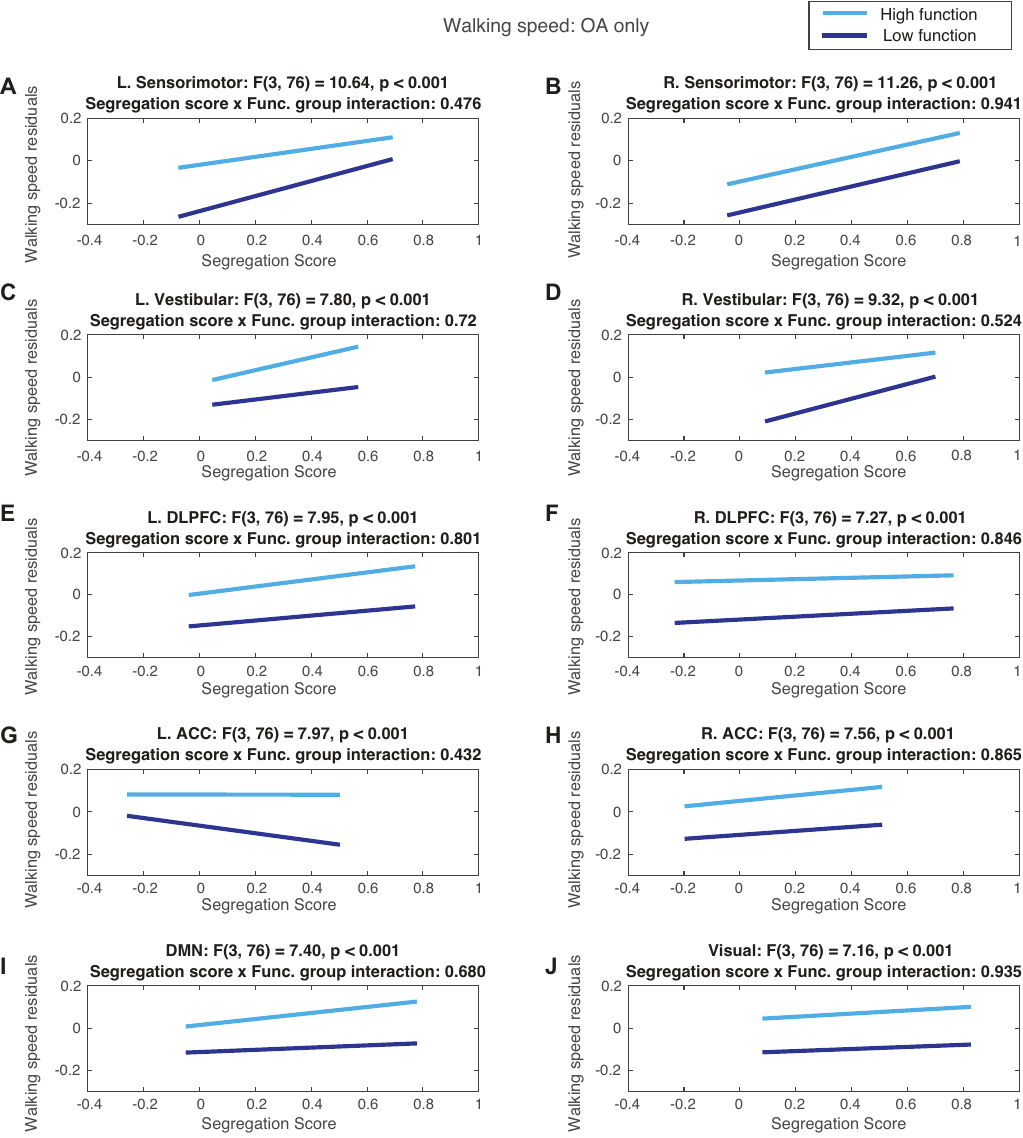

Supplementary figure 3. Univariate linear regression between walking speed corrected for biological sex and segregation scores for each network, with physical function groups. Younger adults were excluded. Lines represent best-fit line. DLPFC = Dorsolateral prefrontal cortex; ACC = Anterior cingulate cortex; DMN = Default mode network.

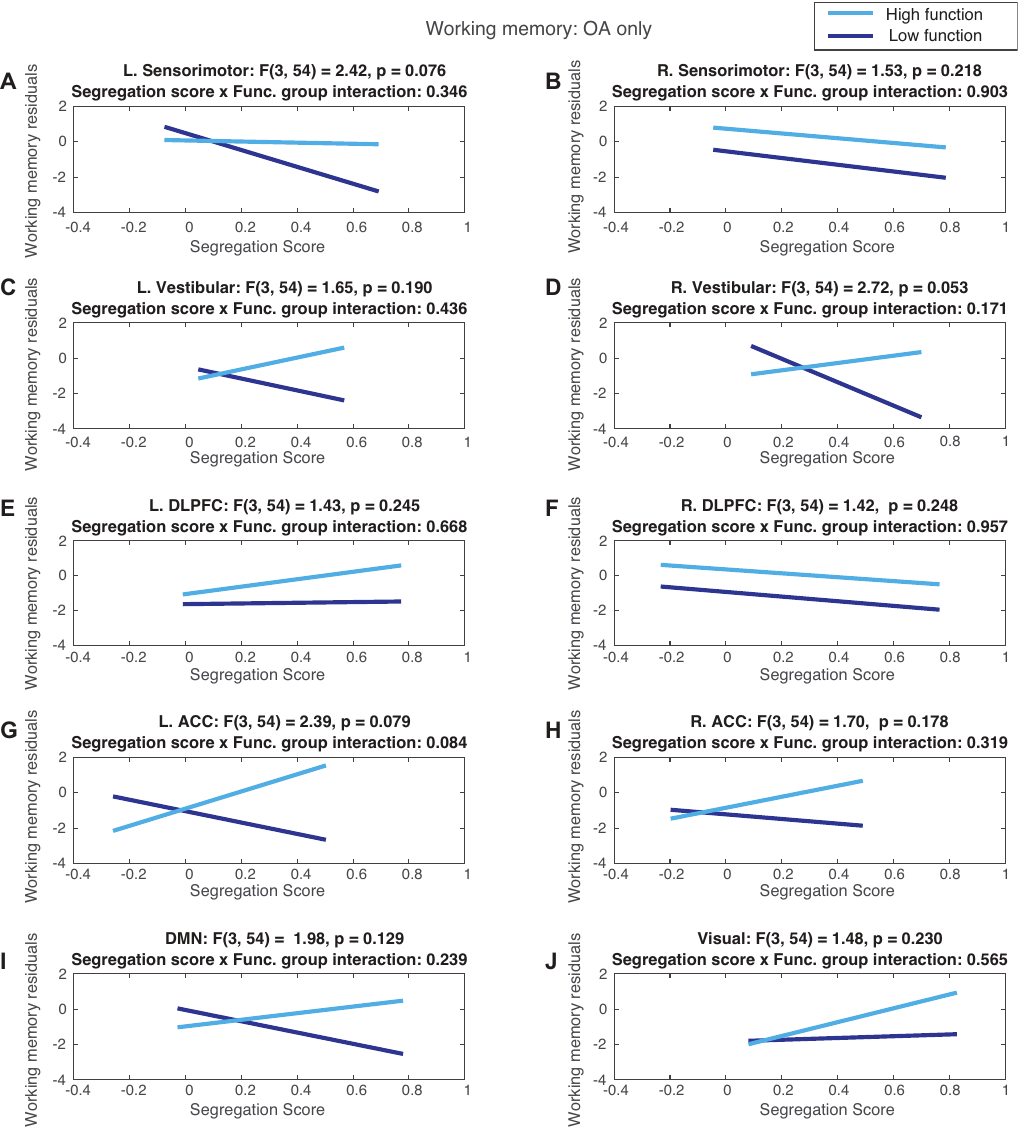

Supplementary figure 4. Univariate linear regression between working memory corrected for biological sex and segregation scores for each network, with physical function groups. Younger adults were excluded. Lines represent best-fit line. DLPFC = Dorsolateral prefrontal cortex; ACC = Anterior cingulate cortex; DMN = Default mode network.
